# Predisposed and learned preferences for multipoint visual statistics in visually naïve newly hatched chicks

**DOI:** 10.1101/2025.06.16.659971

**Authors:** Mirko Zanon, Bastien S. Lemaire, Eugenio Piasini, Riccardo Caramellino, Caroline Nallet, Vijay Balasubramanian, Judit Gervain, Davide Zoccolan, Giorgio Vallortigara

## Abstract

**Significance statement:** We show that visually naïve chicks spontaneously prefer specific multipoint correlation patterns, mirroring preferences seen in humans and rats, and reflecting the most informative structures in natural scenes. This provides evidence that efficient coding mechanisms may be innately driven by evolutionary predispositions. Notably, early visual experience through imprinting can alter these preferences, highlighting a role for learning in shaping visual processing.

Recent studies have revealed that human and non-human animals (rats) can detect luminance distribution and correlations between pixels in an image (ranging from 2-point to 4-point). This sensitivity is believed to stem from optimization processes in the visual system that operate through efficient coding mechanisms to extract the most informative image statistics from the environment. However, it is yet to be determined whether this optimization is evolutionarily given by inborn mechanisms or shaped by visual experience. Here we report that newly-hatched visually naïve domestic chicks spontaneously prefer to approach luminance, 2-point and 4-point correlation patterns (respectively, horizontal lines and rectangular patterns), while showing no preference for 3-point correlation over white noise controls. This parallels the ranking observed in adult humans and rats, thus suggesting that evolutionarily given biological predispositions largely drive efficient coding of natural images. We also found that learning by exposure to visual stimuli, as occurs naturally during visual imprinting, induced a preference for white noise over point correlation patterns in chicks exposed to 3- and 4-point patterns. We hypothesize that this behavior could reflect chicks’ preference for stimuli of lower statistical complexity.

## Introduction

A prominent theory in the field of systems neuroscience is that sensory systems have evolved to efficiently encode the structure of the natural world. According to the hypothesis of efficient coding, neuronal tuning is shaped by the need to optimally encode the statistical structure of input signals(1–6). In vision, this means that our perception should be specifically tuned to the statistical features that are the most informative about the visual world. This idea raises a fundamental question: how do basic image properties shape visual perception in animals?

Natural images can, for example, be statistically characterized in terms of the correlations between their pixels (e.g., up to the fourth order, correlations between values of four adjacent pixels in a 2 x 2 disposition; see Figure 1a; (7, 8)). Correlated pixels define regions, patterns, and objects. Indeed, while the average of single pixel values describes the overall luminance of an image, 2-point correlations give rise to horizontal, vertical and oblique patterns; 3-point correlations define L-shaped forms; 4-point correlations induce rectangles and squares in an image (Figure 1a). Higher-order correlations would reflect more complex visual textures (not considered in these studies). By contrast, an image with no pixel correlations is usually called ‘white noise’: in such a pattern, the pixels are randomly set as white or black independently of one another and the multi-point correlations are all minimized.

**Figure 1.**
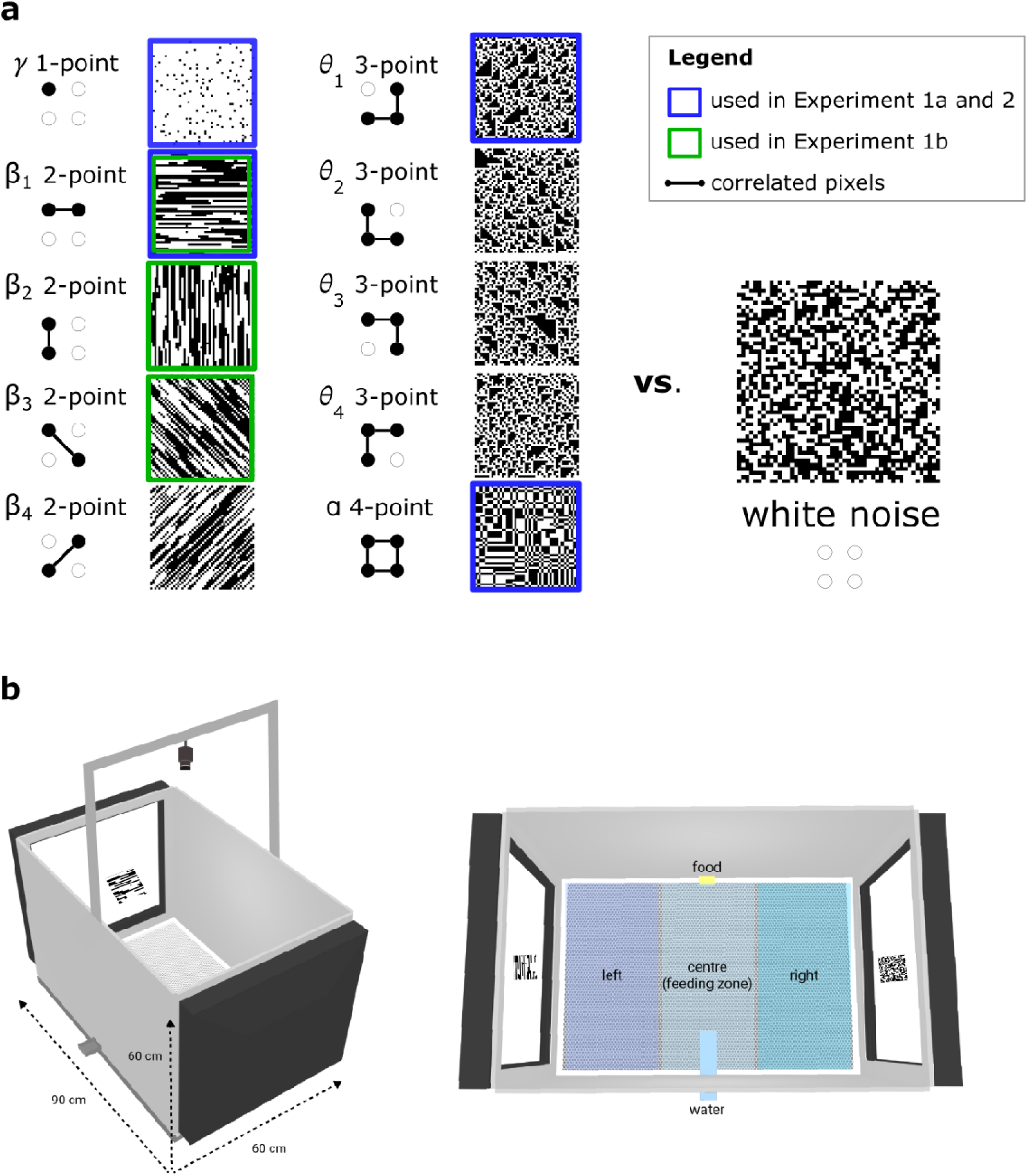
Stimuli and apparatus. ***a)*** Example of textures with defined statistics (correlation between up to four pixels in a 2 x 2 square arrangement as depicted in the gliders on the left of the patterns, with black dots and lines indicating the involved pixel correlation arrangement for the specific statistics). Each independent coordinate (L, β*_1_,* β*_2_,* β*_3_,* β*_4_,* L*_1_,* L*_2_,* L*_3_,* L*_4_,* L as reported in Victor and Conte (8) describes a given correlation pattern (respectively 1-point, 2-point horizontal, 2-point vertical, 2-point oblique to left, 2-point oblique to right, 3-point corner bottom right, 3-point corner bottom left, 3-point corner top right, 3-point corner top left, 4-point correlation). White noise images have all pixel correlations (up to 4^th^ order) equal to zero. See Victor and Conte (8) for a detailed theoretical background. The textures highlighted with a green contour are the ones used in Experiment 1a and 2, while the textures with a blue contour are used in Experiment 1b. ***b)*** Schematic illustration of the experimental setup (9). Lateral view and cage dimensions (left), and upper view with the different choosing areas depicted in different colors (right).

By analyzing these features in natural scenes, the authors of (7, 8) reported that, across natural image patches, there is a prominent variation (bigger variance in the frequency of appearance) of 2-point correlations, followed by 4-point and finally 3-point correlations.

According to the hypothesis of efficient neural coding, if our sensory systems encode environmental signals in an information theoretically optimal way, then these natural image statistics should be reflected in the sensitivities of our visual system: we should be better at perceiving the more variable, i.e. more informative and thus more salient, statistical structures in natural scenes.

To test this idea, Hermundstad et al. (7) conducted psychophysical experiments on human participants, measuring sensitivity to these different visual textures in a discrimination task. Participants were asked to discriminate between the textures, and their perceptual sensitivity was compared to predictions based on the statistics of natural images discussed above. The results showed that human sensitivity was higher for more variable multi-point correlations, supporting the idea that the brain prioritizes processing unpredictable features. In particular, participants performed best with the 2-point, followed by 4-point and lastly 3-point patterns. Such larger sensitivity to the most salient features in natural scenes provides a strong indication that our nervous system encodes visual information efficiently, i.e., it extracts the most information at the lowest possible cost by giving more importance to more informative (i.e. less predictable) features.

Intriguingly, these findings have been recently duplicated in rats (10). After preliminary training where rats learned to differentiate white noise from structured textures (correlation patterns), the animals were tested in a behavioral discrimination task. Their performance was measured by presenting stimuli with varying levels of correlation and fitting psychometric curves. The results revealed that rats, like humans, exhibit a clear sensitivity ranking (better with 1-or 2-point than with 4- and 3-point correlations), supporting the idea that visual perception follows efficient coding principles.

Whether the same sensitivity ranking can be observed in non-mammalian vertebrates is currently unknown. The efficient neural coding hypothesis makes the strong prediction that this should be the case, since any visual species should effectively allocate computational resources to encode the most informative features in natural images. This expectation is reinforced by the similarities of the visual systems among vertebrates (11–13).

The origins of efficient coding mechanisms remain largely unexplored, i.e. whether they reflect learning during ontogeny (e.g., by exposure to natural image statistics, encoded directly in an individual nervous system; (2, 14–16)) or evolutionarily-given inborn predispositions (e.g., by exposure to natural image patterns during natural history, encoded in the genome; (17, 18)) or a mix of both.

To address these questions, we studied perception of multipoint correlation patterns in newly-hatched domestic chicks, a precocial species ideal for the investigation of biological predispositions and early learning (18–24). Newly-hatched chicks possess a range of predisposed visual preferences that guide early learning for pecking at specific food colours and shapes (25, 26), detection of animacy (20), to be used in recognition of social partners (27–29), prey (30, 31) and predators (32). We focus here on spontaneous (innate) preferences for visual textures defined by multipoint correlations and on how experience with these visual stimuli could shape chicks’ choices by filial imprinting, a form of exposure learning.

In Experiment 1, chicks were hatched in the dark to prevent any exposure to statistical regularities of the visual environment and then tested in free-choice tests with specified 1-point statistics (luminance), 2-point correlations (horizontal lines), 3-point correlations (L-shapes with corner to the bottom right) and 4-point correlations (rectangular patterns) *vs.* white noise (Experiment 1a; see Figure 1a). We also tested chicks with additional 2-point patterns not used in previous animal studies, which produced horizontal, vertical and oblique line patterns (Experiment 1b; see Figure 1a), to provide data on possible orientational biases during exposure to asymmetric and not rotationally invariant configurations and to allow for a better comparison with human studies (7).

In Experiment 2, we investigated whether exposure to different correlation patterns affects learning by filial imprinting. To this end, we exposed newly-hatched visually naïve chicks to the different texture statistics (used in Experiment 1a) for one day and then tested their preferences for the familiar (imprinted) texture *vs.* white noise (and vice versa, with imprinting on the noise pattern).

### Experiment 1: Spontaneous choice

In this experiment, we assessed the innate preference (and discriminability) of chicks for different texture statistics over white noise with free-dual-choice tests. The number of males and females was balanced in each group (see Figure 2 and Figure 3 for the relative numbers).

**Figure 2.**
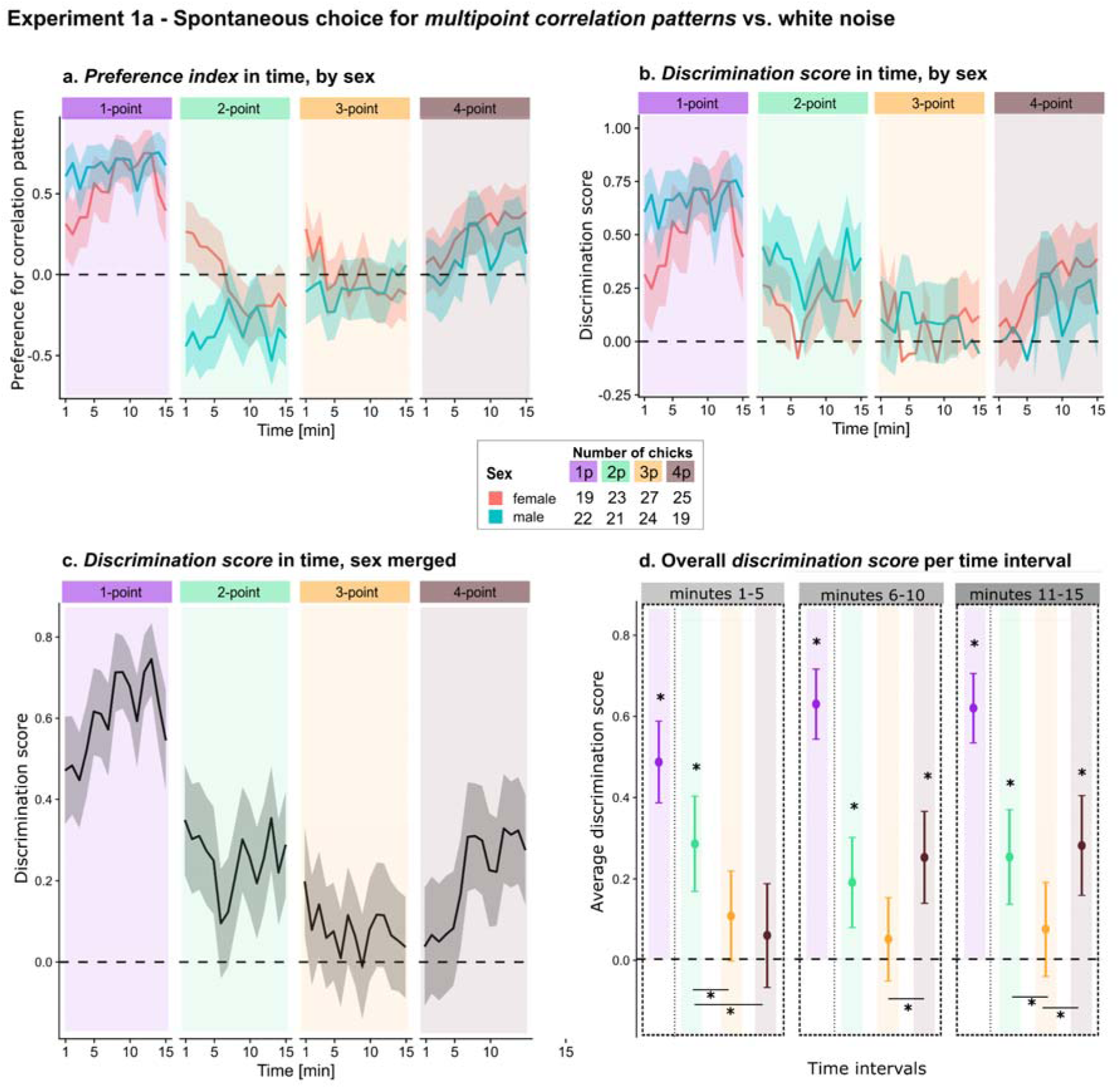
Results for Experiment 1a. ***a)*** Preference index (>0: preference for the correlation pattern) during the first 15 minutes following the first choice, divided by conditions and sex. ***b)*** Discrimination score (>0: preference for pattern or noise) during the first 15 minutes following the first choice, divided by conditions and sex. ***c)*** Discrimination score (preference for pattern or noise) during the first 15 minutes following the first choice, divided by conditions, with sex merged. ***d)*** Average discrimination score in time intervals of 5 minutes (respectively average across minutes 1-5, minutes 6-10, and minutes 11-15). Dotted horizontal lines indicate the chance level (i.e., no preference or discrimination); asterisks (*) indicate statistical significance against the chance level (p < 0.05).

**Figure 3.**
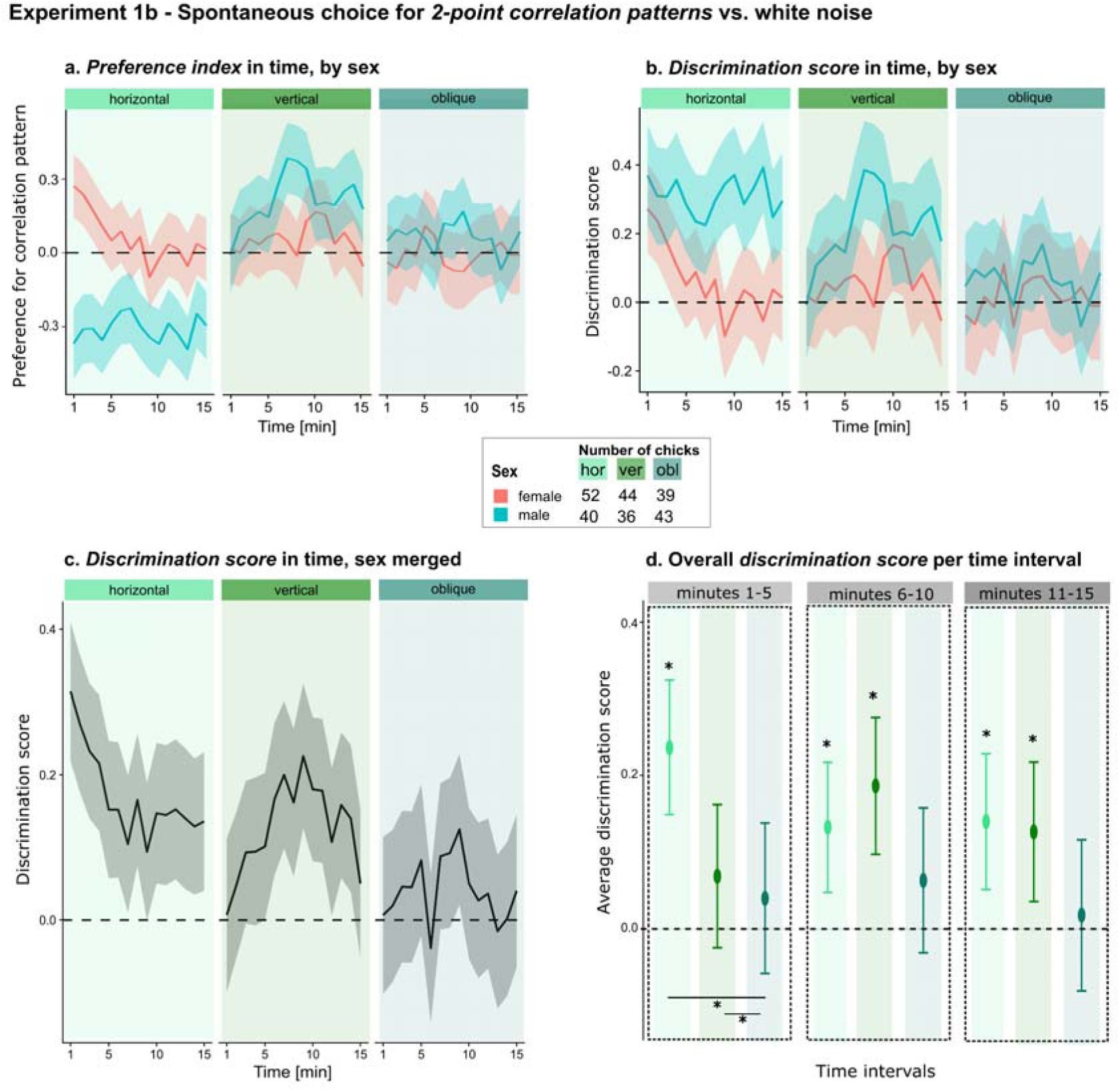
Results for Experiment 1b. ***a)*** Preference index (>0: preference for the correlation pattern) during the first 15 minutes following the first choice, divided by conditions and sex. ***b)*** Discrimination score (>0: preference for pattern or noise) during the first 15 minutes following the first choice, divided by conditions and sex. ***c)*** Discrimination score (preference for pattern or noise) during the first 15 minutes following the first choice, divided by conditions, with sex merged. ***d)*** Average discrimination score in time intervals of 5 minutes (respectively average across minutes 1-5, minutes 6-10, and minutes 11-15). Dotted horizontal lines indicate the chance level (i.e., no preference or discrimination); asterisks (*) indicate statistical significance against the chance level (p < 0.05).

In Experiment 1a we tested the same four texture statistics that (10) studied in rats: 1-point, 2-point horizontal, 3-point, 4-point correlations.

In Experiment 1b we tested different kinds of 2-point patterns (yielding bands with different orientations: horizontal, vertical and oblique) to better investigate possible orientational biases during the exposure to asymmetric and not rotationally invariant configurations. This was motivated by the fact that horizontal 2-point patterns align with the stimulus motion direction, while vertical ones are perpendicular, potentially causing illusions like the Barber-Pole effect (33, 34). Moreover, chicks tend to prefer objects moving along their elongation axis, a possible cue for animacy (35). In addition, this allowed a finer comparison with the sensitivity of humans, who have been found to be more sensitive to cardinal (horizontal, vertical) than oblique orientations (7, 36).

## Results

### Experiment 1a

Looking at the chicks’ preference over time (Figure 2a), it is clear that males and females show similar time-varying trends for 1-, 3- and 4-point patterns. For example, both sexes show a strong preference for the 1-point luminance pattern as compared to white noise (e.g., females, minute 5: PI = 0.56 ± 0.17, 95% C.I. [0.20, 0.93]; females, minute 10: PI = 0.65 ± 0.15, 95% C.I. [0.32, 0.97]; males, minute 5: PI = 0.66 ± 0.14, 95% C.I. [0.38, 0.95]; males, minute 10: PI = 0.71 ± 0.11, 95% C.I. [0.47, 0.94]). For the 3-point patterns chicks’ behavior is always at chance (e.g., females, minute 5: PI = −0.06 ± 0.16, 95% C.I. [−0.40, 0.28]; females, minute 10: PI = −0.08 ± 0.17, 95% C.I. [−0.43, 0.28]; males, minute 5: PI = −0.23 ± 0.17, 95% C.I. [−0.59, 0.13]; males, minute 10: PI = −0.08 ± 0.19, 95% C.I. [−0.47, 0.30]). For the 4-point patterns, preference towards the correlation pattern emerges over time (e.g., females, minute 1: PI = 0.07 ± 0.19, 95% C.I. [−0.33, 0.47]; females, minute 15: PI = 0.39 ± 0.18, 95% C.I. [0.02, 0.75]; males, minute 1: PI = 0.00 ± 0.23, 95% C.I. [−0.47, 0.48]; males, minute 15: PI = 0.13 ± 0.21, 95% C.I. [−0.31, 0.57]). For 2-point patterns, the two sexes show opposite preferences: males show a stable preference for white noise, while females spend more time close to the multipoint pattern during the first minutes of test (e.g., females, minute 1: PI = 0.26 ± 0.19, 95% C.I. [−0.13, 0.66]; females, minute 15: PI = −0.39 ± 0.18, 95% C.I. [−0.76, −0.02]; males, minute 1: PI = −0.44 ± 0.20, 95% C.I. [−0.85, −0.03]; males, minute 15: PI = −0.39 ± 0.18, 95% C.I. [−0.76, −0.02]). The different initial preference for the two sexes in the 2-point group is also visible from the first choice (Supplementary Figure 1). All data for the minute-by-minute choice can be found in Supplementary Table 1.

If we compute the discrimination score (strength of choice, independent of its direction), the curves for males and females align in their trend for all conditions (Figure 2b), as confirmed by permutation test comparisons over choices during the 15 minutes of testing (1-point, male vs. female: p=0.47; 2-point, male vs. female: p=0.30; 3-point, male vs. female: p=0.97; 4-point, male vs. female: p=0.48). Data merged across sexes are shown in Figure 2c.

Disentangling the complex time profiles by averaging the score in 5-minute time bins (Figure 2d), we found a stable discrimination in time for 1-point (average discrimination score for minutes 1-5, permutation test against chance p<0.01; average discrimination score for minutes 6-10, permutation test against chance p<0.01; average discrimination score for minutes 11-15, permutation test against chance p<0.01), as well as for 2-point patterns (average discrimination score for minutes 1-5, permutation test against chance p<0.01; average discrimination score for minutes 6-10, permutation test against chance p<0.01; average discrimination score for minutes 11-15, permutation test against chance p<0.01). The 4-point pattern was discriminated from the middle of the 15 minutes (average discrimination score for minutes 1-5, permutation test against chance p>0.05; average discrimination score for minutes 6-10, permutation t-test against chance p<0.01; average discrimination score for minutes 11-15, permutation test against chance p<0.01). By contrast, the 3-point pattern was never discriminated in any time interval (average discrimination score for minutes 1-5, permutation t-test against chance p>0.05; average discrimination score for minutes 6-10, permutation test against chance p>0.05; average discrimination score for minutes 11-15, permutation t-test against chance p>0.05).

Comparisons between the actual multipoint correlation patterns (2-point, 3-point and 4-point conditions; 1-point is not included in this comparison since it has a different saliency being related to luminance) show differences between 2-point vs. 3-point (permutation Holm corrected p=0.04), and 2-point vs. 4-point (permutation Holm corrected p=0.02) in the first time interval (minutes 1-5); between 4-point vs. 3-point (permutation Holm corrected p=0.03) in the second time interval (minutes 6-10); and between 2-point vs. 3-point (permutation Holm corrected p=0.04), and 4-point vs. 3-point (permutation Holm corrected p=0.02) in the last time interval (minutes 11-15).

### Experiment 1b

Looking at the chicks’ preference over time (Figure 3a), it is clear that males and females show similar time-varying trends for 2-point vertical and oblique patterns. For example in the vertical group both sexes show a slight preference for the pattern (e.g., females, minute 5: PI = 0.07 ± 0.14, 95% C.I. [−0.21, 0.35]; females, minute 10: PI = 0.17 ± 0.13, 95% C.I. [−0.10, 0.44]; males, minute 5: PI = 0.15 ± 0.15, 95% C.I. [−0.16, 0.45]; males, minute 10: PI = 0.20 ± 0.15, 95% C.I. [−0.11, 0.50]). For the oblique group, there is no preference (e.g., females, minute 3: PI = 0.02 ± 0.15, 95% C.I. [−0.28, 0.31]; females, minute 12: PI = 0.01 ± 0.15, 95% C.I. [−0.29, 0.31]; males, minute 3: PI = 0.07 ± 0.15, 95% C.I. [−0.23, 0.37]; males, minute 12: PI = 0.06 ± 0.15, 95% C.I. [−0.23, 0.35]). For 2-point horizontal patterns,the two sexes initially show opposite preferences: males show a stable preference for white noise, while females spend more time close to the multipoint pattern during the first minutes of test (e.g., females, minute 1: PI = 0.27 ± 0.13, 95% C.I. [0.02, 0.53]; females, minute 15: PI = 0.01 ± 0.13, 95% C.I. [−0.25, 0.27]; males, minute 1: PI = −0.37 ± 0.14, 95% C.I. [−0.66, −0.08]; males, minute 15: PI = −0.30 ± 0.14, 95% C.I. [−0.58, −0.01]). The different initial preference for the two sexes in the horizontal group is also visible from the first choice (Supplementary Figure 2). All data for the minute-by-minute choice can be found in Supplementary Table 2.

By computing the discrimination score (strength of choice, independent of its direction), the curves for males and females align in their trend for all conditions (Figure 3b), as confirmed by permutation test comparisons (horizontal, male vs. female: p=0.1; vertical, male vs. female: p=0.3; oblique, male vs. female: p=0.7). Data merged across sexes are shown in Figure 4c.

**Figure 4.**
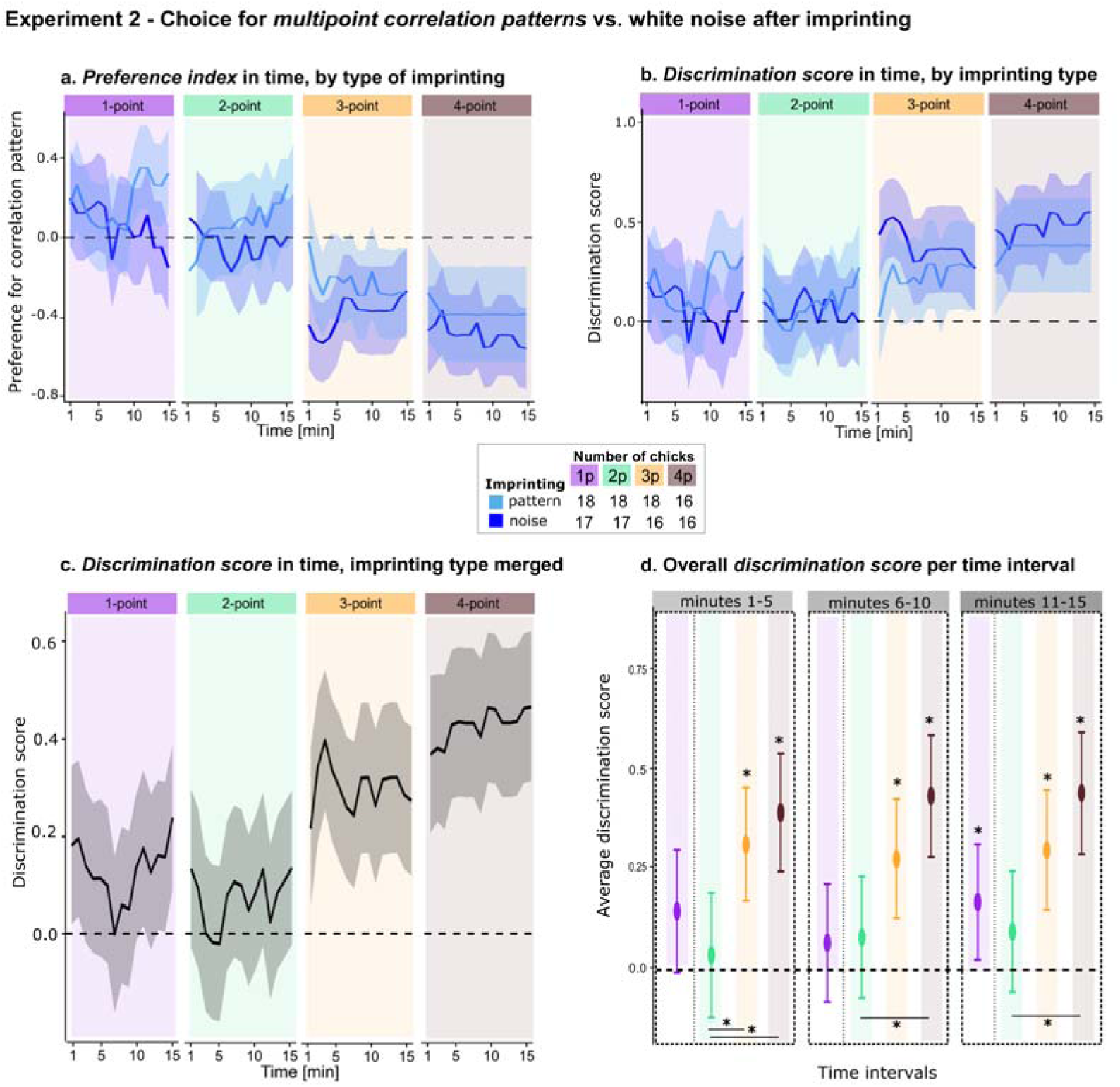
Results for Experiment 2. ***a)*** Preference index (>0: preference for the correlation pattern) during the first 15 minutes following the first choice, divided by conditions and type of imprinting. ***b)*** Discrimination score (>0: preference for pattern or noise) during the first 15 minutes following the first choice, divided by conditions and type of imprinting. ***c)*** Discrimination score (preference for pattern or noise) during the first 15 minutes following the first choice, divided by conditions, with type of imprinting merged. ***d)*** Average discrimination score in time intervals of 5 minutes (respectively average across minutes 1 to 5, minutes 6 to 10, and minutes 11 to 15). Dotted horizontal lines indicate the chance level (i.e., no preference or discrimination); asterisks (*) indicate statistical significance against the chance level (p < 0.05).

Disentangling the complex time profiles by averaging the score in 5-minute time bins (Figure 3d), we found a stable discrimination in time for the 2-point horizontal pattern (average discrimination score for minutes 1-5, permutation test against chance p<0.01; average discrimination score for minutes 6-10, permutation test against chance p<0.01; average discrimination score for minutes 11-15, permutation test against chance p<0.01). The vertical pattern was discriminated towards the middle-end of the 15 minutes (average discrimination score for minutes 1-5, permutation test against chance p>0.05; average discrimination score for minutes 6-10, permutation test against chance p<0.01; average discrimination score for minutes 11-15, permutation test against chance p<0.01). By contrast, the oblique pattern was never discriminated at any time interval (average discrimination score for minutes 1-5, permutation test against chance p>0.05; average discrimination score for minutes 6-10, permutation test against chance p>0.05; average discrimination score for minutes 11-15, permutation test against chance p>0.05).

Comparisons between the various 2-point correlation patterns show differences between horizontal vs. oblique (permutation Holm corrected p<0.01), and horizontal vs. vertical (permutation Holm corrected p=0.01) patterns in the first time interval (minutes 1-5).

## Discussion

### Experiment 1a

We found that chicks spontaneously preferred 1-point luminance textures and 2-point correlation textures. Moreover, a preference for 4-point textures was mainly observed later during the time course of the test (minutes 6-15). While in general the choice is consistent across sexes, 2-point patterns are processed differently in the first minutes (see Figure 2a and first choice, Supplementary Figure 1): females have an initial preference for the 2-point pattern, while males have a stable preference for the white noise patterns over the multipoint statistics. No preference at all was found for 3-point patterns.

1-point patterns define changes in the luminance of an image, a cue that chicks can easily use for discrimination (37). Here, we found a spontaneous preference for brighter stimuli, which is in line with previous findings in domestic chicks (38, 39).

Interestingly, for the 2-point horizontal patterns males and females expressed different initial preferences, with males preferring white noise and females the correlation pattern. The reason for this sex difference is unclear, though a variety of early visual processing mechanisms, from biological motion to responses to face-like stimuli seem to be modulated by sex in young chicks (17, 19, 21, 40, 41).

Still, irrespective of its direction, the spontaneous discrimination of image statistics seems to follow a ranking that mirrors the one documented in humans and rats (7, 10). For rats the 1-point and 2-point patterns are most discriminable from white noise, followed by the 4-, and finally the 3-point patterns (10). Similarly, chicks easily discriminate 1-point patterns from white noise, and they show discrimination for 2-point horizontal patterns and for 4-point textures; while no discrimination occurs at all for 3-point patterns. This ranking is also partially consistent with the one reported by (7), although, unlike humans, chicks do not show any preference for oblique 2-point textures.

In interpreting these findings, it is important to consider the strengths and limitations of our spontaneous preference tests for measuring visual perception. Finding a significant preference for a given texture *vs.* white noise means that the two patterns are discriminable by the chicks’ visual system. By contrast, the lack of a preference does not necessarily imply that the texture is not perceived as different from white noise. Chicks could still be able to discriminate between the two stimuli, without showing any tendency to stay closer to one than to the other. Similarly, the strength of the preference could reflect both the perceptual discriminability of the visual patterns and the innate level of attractiveness of one stimulus over the other.

These caveats notwithstanding, our results seem generally consistent with the hypothesis of efficient coding mechanisms, i.e., that animals’ visual systems are adapted to collect the most informative features from natural scenes, since the observed ranking largely matches the variability of the statistics in natural images. Our results further suggest that these mechanisms are already available at birth, since they are based on spontaneous preferences in visually naïve animals. This, of course, does not exclude the possibility that evolutionarily-driven predispositions could be modulated by exposure to the actual features of a natural environment during development (7, 10).

### Experiment 1b

In this experiment, we found consistent results with Experiment 1a for the horizontal group, thus replicating those findings. Males and females had opposite initial spontaneous preferences when exposed to the horizontal 2-point correlation patterns against white noise. However, interestingly, the spontaneous preference and the sex differences faded away when the line orientations were changed (vertical and oblique cases; Figure 3). Still, for the vertical pattern there is a delayed discrimination (at minutes 6-15), while the oblique pattern was not discriminated (or preferred to white noise) at any time interval. These findings suggest that chicks do not only focus on the mere statistical 2-point correlations present in images but also use other information when showing a spontaneous preference. In our experimental setup, images were moving on the horizontal axis of the screen to increase their attractiveness (moving stimuli have been described to be more attractive than stationary ones in filial imprinting; (27, 42–44)). When the 2-point pattern is horizontally oriented, the lines are parallel to the motion direction, while they are perpendicular in the case of vertical patterns. This different relation between the pattern direction and the direction of motion could cause some sort of illusion in specific configurations, as known, *e.g.*, for the Barber-Pole illusion (33, 34). Moreover, it was shown that chicks prefer objects moving along their main axis of elongation, since this can be recognized as a sign of animacy (35); see (43) for a review). This could account for the effects of orientation in the 2-point pattern groups, though it remains unclear why motion perception direction should be modulated by sex.

### Experiment 2: Discrimination after imprinting

In this experiment, we tested the learned preferences of chicks imprinted on different kinds of correlation statistics or on white noise. Four different groups of chicks were imprinted on textures defined by different multipoint correlations (respectively, with 1-, horizontal 2-, 3- and 4-point) and later tested in a free-dual-choice test between the imprinted texture and white noise. Four other independent groups of chicks were imprinted on white noise images and then tested with white noise *vs.* one of the four texture statistics. The white noise stimuli used for the four groups were equivalent.

## Results

Chicks imprinted above chance with all four patterns and white noise, meaning they spent most of their time close to the imprinting stimulus during the 14 hours of exposition (see Supplementary Figure 4). Averaging across this exposition period, chicks spent 65% of their time (95% C.I. [62,69%], Cohen’s d = 0.77) close to the displayed object.

Concerning the dual choice test phase following imprinting, examination of the chicks’ preference in time (Figure 4a and 4b) makes it clear that chicks imprinted on both the multipoint patterns and white noise show similar time-varying trends at test. For example, for the 1-point statistic both groups show a slight preference for the luminance pattern over noise (e.g., imprinted on noise, minute 1: PI = 0.20 ± 0.24, 95% C.I. [−0.31, 0.71]; imprinted on pattern, minute 1: PI = 0.17 ± 0.23, 95% C.I. [−0.31, 0.65]). For 2-point patterns the preference is close to chance (e.g., imprinted on noise, minute 1: PI = 0.10 ± 0.24, 95% C.I. [−0.41, 0.61]; imprinted on pattern, minute 1: PI = −0.17 ± 0.23, 95% C.I. [−0.65, 0.31]). For 3- and 4-point patterns the preference is stable towards the white noise (e.g., 3-point imprinted on noise, minute 2: PI = −0.51 ± 0.20, 95% C.I. [−0.93, −0.08], minute 14: PI = −0.30 ± 0.22, 95% C.I. [−0.76, 0.16]; 3-point imprinted on pattern, minute 2: PI = −0.19 ± 0.23, 95% C.I. [−0.67, 0.28], minute 14: PI = −0.27 ± 0.22, 95% C.I. [−0.74, 0.19]; 4-point imprinted on noise, minute 2: PI = −0.44 ± 0.20, 95% C.I. [−0.86, −0.01], minute 14: PI = −0.54 ± 0.20, 95% C.I. [−0.97, −0.12]; 4-point imprinted on pattern, minute 2: PI = −0.33 ± 0.23, 95% C.I. [−0.82, 0.17], minute 14: PI = −0.38 ± 0.24, 95% C.I. [−0.89, 0.12]). Choices at all time points are reported in Supplementary Table 3.

The similarity, at test, in behaviour between groups imprinted with a pattern or noise is also visible from first choices (Supplementary Figure 5), and is confirmed by permutation test comparisons on the discrimination score (1-point, imprinting with pattern vs. noise: permutation p=0.6; 2-point, imprinting with pattern vs. noise: permutation p=0.7; 3-point, imprinting with pattern vs. noise: permutation p=0.6; 4-point, imprinting with pattern vs. noise: permutation p=0.7). Data merged across types of imprinting are shown in Figure 4c.

Simplifying the time profiles by averaging the score in 5 minutes time bins (Figure 4d), we found a stable discrimination in time for 3-point (average discrimination score for minutes 1-5, permutation test against chance p<0.01; average discrimination score for minutes 6-10, permutation test against chance p<0.01; average discrimination score for minutes 11-15, permutation test against chance p<0.01), as well as for 4-point patterns (average discrimination score for minutes 1-5, permutation test against chance p<0.01; average discrimination score for minutes 6-10, permutation test against chance p<0.01; average discrimination score for minutes 11-15, permutation test against chance p<0.01). The 1-point pattern was discriminated towards the end of the 15 minutes (average discrimination score for minutes 1-5, permutation test against chance p>0.05; average discrimination score for minutes 6-10, permutation test against chance p>0.05; average discrimination score for minutes 11-15, permutation test against chance p=0.03). By contrast, the 2-point pattern was never discriminated at any time interval (average discrimination score for minutes 1-5, permutation test against chance p>0.05; average discrimination score for minutes 6-10, permutation test against chance p>0.05; average discrimination score for minutes 11-15, permutation test against chance p>0.05).

Comparisons between the multipoint correlation patterns (2-point, 3-point and 4-point conditions; 1-point is not included in this comparison since it has a different saliency being related to luminance) show differences between 2-point vs. 3-point patterns (permutation Holm corrected p<0.01), and 2-point vs. 4-point patterns (permutation Holm corrected p<0.01) in the first time interval (minutes 1-5); between 4-point vs. 2-point patterns (permutation Holm corrected p<0.01) in the second time interval (minutes 6-10); and between 2-point vs. 4-point patterns (permutation Holm corrected p<0.01) in the last time interval (minutes 11-15).

## Discussion

Previous studies investigated sensitivity to different kinds of correlation statistics using operant conditioning procedures, in which subjects were required to explicitly discriminate structured textures from white noise patterns (7, 10). In the case of rats, this was achieved by teaching them an association between stimulus identity and reward ports. In Experiment 2, we explored whether pixel correlations could affect a different, non-associative form of learning, namely filial imprinting (there is evidence that imprinting attains more to declarative-like rather than procedural forms of learning; (45)).

Our results suggest that chicks imprint equally well on all patterns and white noise, spending 65% of their time (above chance) close to the displayed object. Still, it is interesting to notice how in (28), chicks underwent a similar procedure with different stimuli, spending 96% of their time close to their imprinting objects. Such a difference suggests that our correlation patterns are probably not the most optimal stimuli for imprinting, possibly also because of the absence of colour.

At test, the 2-point groups showed no preference, as if chicks did not differentiate between the correlation patterns and white noise. The preference for 1-point luminance appears only towards the end of the test (minutes 11-15). However, we know from the results of Experiment 1 that chicks can discriminate these two patterns. Thus, it appears that, after imprinting, the difference of 2-point patterns (and to some extent 1-point patterns) from white noise was no longer judged relevant enough to elicit a strong differential behavioral approach. Interestingly, however, stronger preferences appeared in the 4- and 3-point patterns. In both these groups, chicks preferred white noise when imprinted on it but also when imprinted on the correlation patterns. This suggests that chicks imprinted with 3- and 4-point patterns tend to explore novel stimuli (the white noise) while chicks imprinted with noise mostly remain attracted to this familiar pattern. Generally speaking, in these cases, white noise seems visually more attractive than 3- and 4-point patterns.

Interestingly, imprinting leads to an outcome that is mirror-reversed with respect to the spontaneous preferences test. Several studies showed that young chicks tend to behave differently in imprinting tests on the basis of the perceived novelty of the stimulus (46–48). Strikingly, completely unfamiliar stimuli tend to be avoided whereas slight novelty is explored (49). Here, the results suggest that 3- and 4-point patterns elicited clear choices, whereas 2- and 1-point did not. The different orders of the correlation patterns show increasing complexity, starting from single-pixel luminance to multiple-pixel correlations. In this sense, white noise can be considered the lowest in complexity, lacking any correlation. Our stimuli thus show increasing complexity starting from noise to 1-, 2-, 3- and 4-point patterns, defined by the number of units involved, as visible by the sketches in Figure 1a), with white noise at the lowest bound. If chicks possess a mechanism to detect pixel correlations, as apparent from the results of Experiment 1, a white noise image can be considered to be closer to e.g. a 2-point than to a 4-point pattern. Our results converge with chicks being sensitive to such an ordering: clear choice in imprinting is apparent when white noise is compared with 3- or 4-point patterns, but not with ‘simpler’ 1- or 2-point patterns.

### Conclusions

We found that newly-hatched visually naïve domestic chicks show a spontaneous preference to approach 1-point, 2-point and 4-point correlation patterns (respectively, luminance, horizontal lines and rectangular patterns), but no preference at all for 3-point correlation pattern versus white noise. This ranking resembles the one observed in adult humans and rats and could suggest evolutionarily-set biological predispositions underlying efficient coding of natural images. Interestingly, we also found that imprinting to visual patterns (both multipoint correlation or noise) induces a stronger preference (and thus a discrimination) for white noise over point correlation patterns in animals exposed to 3- and 4-point stimuli, while the preference for 2- and 1-point patterns is no longer present at the beginning of the test. This behavior could reflect an attraction to stimuli of lower statistical complexity and an avoidance of strongly novel patterns, with a modulation provided by innate preferences.

Our results provide the first evidence for an evolutionary grounding of efficient coding mechanisms and open the way to further investigation of their underlying neural and genetic mechanisms.

## Materials and methods

### Subjects

The number of animals tested in each group was a priori determined by a power analysis considering an effect size (Cohen’s d) of 0.55 in the spontaneous preference test (inferred from Caramellini et al.(10)) and of 0.75 in the imprinting test, inferred from Lemaire et al. (28)), and an alpha of 0.05. Twenty-eight individuals per group for the spontaneous preference test (Experiment 1a) and 16 individuals per group in the imprinting test (Experiment 2) were required to achieve a power of 0.8. Experiment 1b (on different orientations of 2-point patterns) was run a posteriori, and the power analysis was based on a more appropriate effect size from Experiment 1a considering the group tested with horizontal 2-point patterns: for an effect size of 0.37 (see Experiment 1b Results section), to reach a power of 0.8, 47 chicks per group were required. Overall, we tested 671 chicks (263 males) of the strain Ross 308, from which we included 529 chicks (184 males) in the analysis (after excluding non-moving animals; see Data analysis subparagraph). The eggs were obtained from a commercial hatchery (Azienda Agricola Crescenti, Brescia) and were incubated in complete darkness in our laboratory under controlled temperature (37.7 °C) and humidity (40% humidity). Three days before hatching, eggs were moved into opaque black boxes within a hatching chamber at 37.7 °C and 60% of humidity. Soon after hatching, chicks were briefly sexed under dim light (based on sexual dimorphism of the wing feathers) and singly housed in testing cages, positioning them in the center. All procedures received approval from the Ethical Committee of the University of Trento and the Italian Ministry of Health (permit number 53/2020-PR released on 21/01/2020).

### Apparatus

All experiments took place in testing cages of 90×60×60 cm, with water and food available *ad libitum*. At two opposite sides of the cage, two high frame rate (120 Hz) screens displayed the stimuli. Image presentation was automated and controlled by a custom Matlab script which ensured the display of stimuli at precise times and randomized stimulus position on the screens in order to avoid positional bias (9). The behaviour of each chick was continuously recorded using an overhead camera. The experimental setup and methods are described in detail in Zanon et al. (9). See Figure 1b for a schematic representation of the set-up.

### Stimuli

Textures with different multipoint correlations and white noise patterns were generated using Metex (50). This program creates textures in which the probability of occurrence of a given correlation can be controlled systematically, while minimizing the contribution of other multipoint correlations to the structure of the images (this is achieved by sampling images from the maximum-entropy distribution that is compatible with the requested correlation^8^). This way, the pixels in the generated texture are as random as possible, while respecting the desired multipoint correlation constraint. In contrast, white noise images contain no multipoint correlations. The space of possible textures is parameterized by what we will refer to as the intensity of the corresponding statistic, which can take on values between −1 and 1. When the intensity is zero, the texture does not contain any structure and, therefore, it is the same as white noise. When the intensity is close to one of its extreme values (i.e., close to either +1 or −1), the structure dictated by the correlation gives rise to textures with the prominent, characteristic features shown in Figure 1a (see Victor and Conte (8) for the mathematical details on how the space of textures is parameterized).

We used high intensity values (+0.9) for all the statistics we tested; in this way, we ensured that the resulting textures were highly discriminable from white noise. For Experiments 1a and 2 we used 1-point, 2-point horizontal, 3-point corner bottom right, and 4-point; while for Experiment 1b we used 2-point horizontal, vertical and oblique left (see Figure 1a). The remaining patterns (2-point oblique right and 3-point corner top left/right and bottom left) were not investigated for time reasons, also considering the supposed scarce biological relevance and novelty compared to the already considered ones.

Thus, we created 10 images for each of the aforementioned statistics of interest (1-point, 2-point horizontal, vertical and oblique, 3-point corner bottom right and 4-point). In all dual-choice phases of all experiments each chick was displaying one of these patterns against white noise (always generated in a new different random pattern); while in the imprinting phase of Experiment 2, patterns of a given statistic were randomly varied spanning the 10 created patterns: in this way we avoided the possibility that the chick can imprint on the identity of a very specific stimulus, considering that the same pattern was never displayed more than once to the same subject across experimental sessions.

The image size was 300×300 pixels (8.3×8.3 cm) to ensure a minimum angle of view for the whole stimulus (estimated for a maximum distance of 90 cm, from one side of the cage to the opposite screen) of 2.9°, which is clearly perceived by chicks (51). A single pixel had a dimension of 0.16 cm, ensuring a minimum angle of view of 0.11° at a distance of 45 cm. Thus, we could be confident that chicks could visually discriminate the different patterns already from the center of the cage.

The stimuli displayed on the screens were moving (translating) horizontally to attract the chicks’ attention. We used the following parameters of our Matlab script (9): translation amplitude 0.32 corresponding to 42.5 cm spanned and translation frequency 0.11 corresponding to 275 cm/min).

## Experimental pipeline

### Experiment 1

Experiments 1a and 1b consisted of spontaneous dual-choice tests. After positioning in the center of the cage, chicks underwent a dual-choice multipoint pattern *vs.* white noise test (each chick was assigned to one of the four specific textures, see Figure 1a). The whole test lasted 30 minutes (to ensure chicks could initiate movement, after which only the first 15 minutes were analyzed). Chicks that stayed in the center of the apparatus and did not make a choice between the presented stimuli for more than 15 minutes were discarded from analysis. Chicks were exposed to one single texture of a given correlation for the whole duration of the test. The experimental texture was displayed on a random side of the cage, while on the opposite screen, the white noise stimulus was displayed. Both stimuli were horizontally translating on screens to increase attractiveness (27, 42–44).

### Experiment 2

This experiment consists of an imprinting exposure followed by a dual-choice test. After positioning in the center of the cage, chicks underwent one day of imprinting with a specific pattern (24 hours divided into 7 sessions of 2 hours: 14 hours of active imprinting, plus a night resting time of 10 hours with black screens; for a reference to the procedure see also Lemaire et al. (28)). During this imprinting phase, the stimulus was displayed only on one screen (while the opposite screen was black). The active screen changed randomly every session (i.e., every 2h) and a new instance of the same imprinting multipoint correlation pattern (or white noise pattern) was displayed in each session: this prevents chicks from learning and focusing on a specific stimulus or screen during the imprinting phase.

Once the imprinting phase was completed, chicks were repositioned at the center of the cage to avoid any positional bias at the beginning of the test (dual-choice texture *vs.* white noise). The test phase was similar to Experiment 1, with both stimuli horizontally translating on the opposite screens. The statistical textures, and consequently of the white noise, were randomly allocated to specific screens.

In order to limit the number of animals used we decided to employ only one sex in Experiment 2: we selected female chicks as they are usually more attached to social stimuli than males in imprinting tests (52–55). Females also showed a stronger preference for textures against white noise in Experiment 1 (see Figure 2a and 3a).

### Data analysis

For all the experiments, the location of the subjects within the arena was analysed with DeepLabCut (56, 57). The animal was considered close to a screen (stimulus) when it was less than 30 cm from it.

We analyzed the first 15 minutes following the subject’s first choice. We discarded from the analyses chicks that stayed in the center of the apparatus and did not make a choice between the presented stimuli for more than 15 minutes (see Supplementary Table 1 for the exact numbers of chicks used and analysed).

To take into account that chick preferences or discrimination performance could vary during different epochs of the 15 minutes, we ran the statistical analysis independently in 5-minute intervals (‘beginning’: minutes 1-5; ‘middle’: minutes 6-10; and ‘end’: minutes 11-15; see details below).

As metrics for evaluating chick preferences, we recorded the first stimulus approached by each animal, and calculated a *preference index* (at single minute resolution) using the formula:

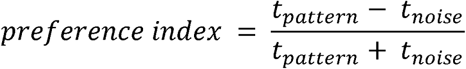

with *t* being the seconds spent close to a stimulus (respectively correlation patterns and white noise). This index reflects the preference of the animals for the displayed stimuli. An index value of +1 indicates that the animals spent all their time close to the multipoint texture; an index value of −1 indicates that the animals spent all their time close to the white noise; while a value of 0 indicates chance level.

We investigated actual choice (between noise and pattern) over time by plotting the chicks’ average *preference index* minute by minute, divided by sex. We took sex into consideration for our analysis of Experiment 1 since a variety of early visual processing mechanisms, from biological motion to responses to face-like stimuli, seem to be modulated by sex in young chicks (17, 19, 21, 40, 41).

To compare results with previous studies (7, 10), we also focused on the general ability to discriminate multipoint patterns from white noise, as opposed to the specific preference for one or the other. To quantify this discrimination, we computed a *discrimination score* based on the *preference index*. For each subgroup, defined by sex, statistical condition, and time interval, we first calculated the group mean of the *preference index*. If this mean was negative, indicating a consistent preference for white noise within the group during that interval, we inverted the sign of the preference indices for all individuals at all time points, in that subgroup and interval. If the mean was positive, no transformation was applied.

This procedure aligns all group-level preferences in the same direction, allowing us to assess the strength of discrimination regardless of stimulus identity. The resulting *discrimination score* thus provides a measure of the magnitude of group-level choices, with higher values reflecting stronger discrimination. Importantly, because the data are still analyzed and plotted across time, individual time points may still show negative values, indicating temporary divergences from the overall (average in time and across subjects) preference trend in the specific interval. We also note that while a positive *discrimination score* reflects a clear pattern vs. noise discrimination, a score of zero does not necessarily indicate an absence of discrimination, but rather may reflect a lack of consistent preference across individuals or of overall (average in time and across subjects) preference as well.

All the statistical analyses were performed on these *discrimination scores* to check if each multipoint pattern was discriminated from noise (i.e., testing *discrimination score* difference from zero) in different time intervals (beginning, middle, and end, as defined above). Analyses were performed in RStudio (version 1.4.17.17) running R (version 4.1.0, R Core Team, 2017).

Specifically, we firstly evaluated the contribution of sex (for Experiment 1) or type of imprinting (for Experiment 2) for each multipoint pattern, by running a permutation test (5000 permutations) on the scores of male vs. female (Experiment 1) or imprinting on noise vs. multipoint pattern (Experiment 2).

If no significant difference between the sexes or type of imprinting was found, the data were merged and the difference of the overall *discrimination score* from chance level zero was evaluated with a permutation test for each time interval. In this case, multiple (5000) random datasets were created by randomly inverting the sign of the preference indices at the single chick level in order to mimic random choice by subjects. In this way, the distribution of the means for the random datasets was constructed. The p-value was thus evaluated as the percentage of random data points more extreme than the real mean, using the formula: p-value = (b+1)/(n_perm+1), where b is the number of values more extreme than the real statistic value, n_perm = 5000 iterations (58).

Finally, to compare differences between multipoint patterns, Holm-corrected permutation tests were used. As in previous works (7), we compared only 2-point, 3-point and 4-point (for Experiment 1a and 2), and 2-point horizontal, vertical and oblique (for Experiment 1b) since the 1-point pattern simply corresponds to luminance variations, and is thus not a multipoint correlation.

## Declarations

### Availability of data and material

All the materials and scripts to replicate the analysis are freely available in the Figshare repository: https://doi.org/10.6084/m9.figshare.28846787.v1

### Authors’ contributions

M.Z., B.S.L. performed the experiments and the overall data analysis. G.V., D.Z. and J.G. supervised the project. M.Z., B.S.L. and E.P. performed statistical analysis. M.Z., B.S.L. and G.V. wrote the initial draft. All the authors contributed to the reshaping, the correction and improvement, and the discussion for the final version of the manuscript.

## Bibliography

1. H. Barlow, Redundancy reduction revisited. Netw. Comput. Neural Syst. 12, 241–253 (2001).

2. G. Matteucci, E. Piasini, D. Zoccolan, Unsupervised learning of mid-level visual representations. Curr. Opin. Neurobiol. 84, 102834 (2024).

3. C. P. Ratliff, B. G. Borghuis, Y.-H. Kao, P. Sterling, V. Balasubramanian, Retina is structured to process an excess of darkness in natural scenes. (2010).

4. E. P. Simoncelli, Vision and the statistics of the visual environment. Curr. Opin. Neurobiol. 13, 144–149 (2003).

5. T. Tesileanu, E. Piasini, V. Balasubramanian, Efficient processing of natural scenes in visual cortex. Front. Cell. Neurosci. 16, 1006703 (2022).

6. G. Tkačik, J. S. Prentice, J. D. Victor, V. Balasubramanian, Local statistics in natural scenes predict the saliency of synthetic textures. Proc. Natl. Acad. Sci. 107, 18149–18154 (2010).

7. A. M. Hermundstad, et al., Variance predicts salience in central sensory processing. eLife 3, e03722 (2014).

8. J. D. Victor, M. M. Conte, Local image statistics: maximum-entropy constructions and perceptual salience. J. Opt. Soc. Am. A 29, 1313 (2012).

9. M. Zanon, B. S. Lemaire, G. Vallortigara, Steps towards a computational ethology: an automatized, interactive setup to investigate filial imprinting and biological predispositions. Biol. Cybern. 115, 575–584 (2021).

10. R. Caramellino, et al., Rat sensitivity to multipoint statistics is predicted by efficient coding of natural scenes. eLife 10, e72081 (2021).

11. G. Vallortigara, “The cognitive chicken: Visual and spatial cognition in a non-mammalian brain” in The Oxford Handbook of Comparative Cognition, (Oxford University Press, 2012), pp. 48–66.

12. E. I. Knudsen, Evolution of neural processing for visual perception in vertebrates. J. Comp. Neurol. 528, 2888–2901 (2020).

13. G. Vallortigara, “Visual cognition and representation in birds and primates” in Vertebrate Comparative Cognition: Are Primates Superior to Non-Primates?, (Kluwer Academic/Plenum Publishers, 2004), pp. 57–94.

14. N. Berardi, T. Pizzorusso, L. Maffei, Critical periods during sensory development. Curr. Opin. Neurobiol. 10, 138–145 (2000).

15. J. S. Espinosa, M. P. Stryker, Development and Plasticity of the Primary Visual Cortex. Neuron 75, 230–249 (2012).

16. G. Matteucci, D. Zoccolan, Unsupervised experience with temporal continuity of the visual environment is causally involved in the development of V1 complex cells. Sci. Adv. 6, eaba3742 (2020).

17. O. Rosa Salva, M. Grassi, E. Lorenzi, L. Regolin, G. Vallortigara, Spontaneous preference for visual cues of animacy in naïve domestic chicks: The case of speed changes. Cognition 157, 49–60 (2016).

18. G. Vallortigara, Born Knowing: Imprinting and the Origins of Knowledge (MIT press, 2021).

19. E. Di Giorgio, et al., Filial responses as predisposed and learned preferences: Early attachment in chicks and babies. Behav. Brain Res. 325, 90–104 (2017).

20. E. Lorenzi, G. Vallortigara, “Evolutionary and Neural Bases of the Sense of Animacy” in The Cambridge Handbook of Animal Cognition, 1st Ed., A. B. Kaufman, J. Call, J. C. Kaufman, Eds. (Cambridge University Press, 2021), pp. 295–321.

21. O. Rosa Salva, U. Mayer, G. Vallortigara, Roots of a social brain: Developmental models of emerging animacy-detection mechanisms. Neurosci. Biobehav. Rev. 50, 150–168 (2015).

22. O. Rosa Salva, et al., Sensitive periods for social development: Interactions between predisposed and learned mechanisms. Cognition 213, 104552 (2021).

23. E. Versace, A. Martinho-Truswell, A. Kacelnik, G. Vallortigara, Priors in Animal and Artificial Intelligence: Where Does Learning Begin? Trends Cogn. Sci. 22, 963–965 (2018).

24. E. Versace, G. Vallortigara, Origins of Knowledge: Insights from Precocial Species. Front. Behav. Neurosci. 9 (2015).

25. G. Gamberale-Stille, B. Tullberg, Fruit or aposematic insect? Context-dependent colour preferences in domestic chicks. Proc. R. Soc. Lond. B Biol. Sci. 268, 2479–2484 (2001).

26. T. J. Roper, N. M. Marples, Colour preferences of domestic chicks in relation to food and water presentation. Appl. Anim. Behav. Sci. 54, 207–213 (1997).

27. J. J. Bolhuis, Mechanisms of avian imprinting: a review. Biol. Rev. 66, 303–345 (1991).

28. B. S. Lemaire, D. Rucco, M. Josserand, G. Vallortigara, E. Versace, Stability and individual variability of social attachment in imprinting. Sci. Rep. 11, 7914 (2021).

29. B. J. McCabe, Visual Imprinting in Birds: Behavior, Models, and Neural Mechanisms. Front. Physiol. 10, 658 (2019).

30. E. Clara, L. Regolin, G. Vallortigara, L. J. Rogers, Chicks prefer to peck at insect-like elongated stimuli moving in a direction orthogonal to their longer axis. Anim. Cogn. 12, 755–765 (2009).

31. E. Mascalzoni, D. Osorio, L. Regolin, G. Vallortigara, Symmetry perception by poultry chicks and its implications for three-dimensional object recognition. Proc. R. Soc. B Biol. Sci. 279, 841–846 (2012).

32. M. Hébert, E. Versace, G. Vallortigara, Inexperienced preys know when to flee or to freeze in front of a threat. Proc. Natl. Acad. Sci. 116, 22918–22920 (2019).

33. R. J. Snowden, T. C. A. Freeman, The visual perception of motion. Curr. Biol. 14, R828–R831 (2004).

34. P. Sun, C. Chubb, G. Sperling, Two mechanisms that determine the Barber-Pole Illusion. Vision Res. 111, 43–54 (2015).

35. O. Rosa Salva, M. Hernik, A. Broseghini, G. Vallortigara, Visually-naïve chicks prefer agents that move as if constrained by a bilateral body-plan. Cognition 173, 106–114 (2018).

36. A. R. Girshick, M. S. Landy, E. P. Simoncelli, Cardinal rules: visual orientation perception reflects knowledge of environmental statistics. Nat. Neurosci. 14, 926–932 (2011).

37. J. F. Zolman, W. J. Lattin, Development of brightness preferences in young chicks: Effects on brightness discrimination learning. J. Comp. Physiol. Psychol. 79, 271–283 (1972).

38. C. H. Clarke, B. R. Jones, Domestic Chicks’ Attraction to Video Images: Effects of Stimulus Movement, Brightness, Colour and Complexity. Int. J. Comp. Psychol. 13 (2000).

39. J. D. Delius, G. Thompson, Brightness Dependence of Colour Preferences in Herring Gull Chicks. Z. Für Tierpsychol. 27, 842–849 (1970).

40. M. Miura, T. Matsushima, Preference for biological motion in domestic chicks: sex-dependent effect of early visual experience. Anim. Cogn. 15, 871–879 (2012).

41. O. Rosa Salva, L. Regolin, G. Vallortigara, Faces are special for newly hatched chicks: evidence for inborn domainLJspecific mechanisms underlying spontaneous preferences for faceLJlike stimuli. Dev. Sci. 13, 565–577 (2010).

42. E. H. Hess, Imprinting. Science 130 (1959).

43. B. S. Lemaire, G. Vallortigara, Life is in motion (through a chick’s eye). Anim. Cogn. 26, 129–140 (2022).

44. K. Z. Lorenz, The Companion in the Bird’s World. The Auk 54, 245–273 (1937).

45. P. Bateson, Is imprinting such a special case? Philos. Trans. R. Soc. Lond. B. Biol. Sci. 329, 125–131 (1990).

46. P. Bateson, Brief exposure to a novel stimulus during imprinting in chicks and its influence on subsequent preferences. Anim. Learn. Behav. 7, 259–262 (1979).

47. P. Bateson, Preferences for familiarity and novelty: A model for the simultaneous development of both. J. Theor. Biol. 41, 249–259 (1973).

48. P. S. Jackson, P. Bateson, Imprinting and exploration of slight novelty in chicks. Nature 251 (1974).

49. E. Versace, M. J. Spierings, M. Caffini, C. Ten Cate, G. Vallortigara, Spontaneous generalization of abstract multimodal patterns in young domestic chicks. Anim. Cogn. 20, 521–529 (2017).

50. E. Piasini, metex - Maximum Entropy TEXtures (1.1.0). (2021). Deposited 2021.

51. C. E. Wisely, et al., The chick eye in vision research: An excellent model for the study of ocular disease. Prog. Retin. Eye Res. 61, 72–97 (2017).

52. M. Cailotto, G. Vallortigara, M. Zanforlin, Sex differences in the response to social stimuli in young chicks. Ethol. Ecol. Evol. 1, 323–327 (1989).

53. G. Vallortigara, Affiliation and Aggression As Related to Gender in Domestic Chicks (Gallus gallus). J. Comp. Psychol. 29, 56–57 (1992).

54. G. Vallortigara, Cailotto, M., Zanforlin, M., Sex differences in social reinstatement motivation of the domestic chick (Gallus gallus) revealed by runway tests with social and nonsocial reinforcement. J. Comp. Psychol. 104, 361–367 (1990).

55. L. Workman, J. Andrew, Simultaneous changes in behaviour and in iateralization during the development of male and female domestic chicks. Anim. Behav. 38 (1989).

56. A. Mathis, et al., DeepLabCut: markerless pose estimation of user-defined body parts with deep learning. Nat. Neurosci. 21, 1281–1289 (2018).

57. T. Nath, et al., Using DeepLabCut for 3D markerless pose estimation across species and behaviors. Nat. Protoc. 14, 2152–2176 (2019).

58. A. C. Davison, D. V. Hinkley, Bootstrap methods and their application (Reprinted with corrections 2003) (1997).

