## Supplementary materials for "Predisposed and learned preferences for multipoint visual statistics in visually naïve newly hatched chicks"

**Content:**

- Supplementary Figures 1-5
- Supplementary Tables 1-3

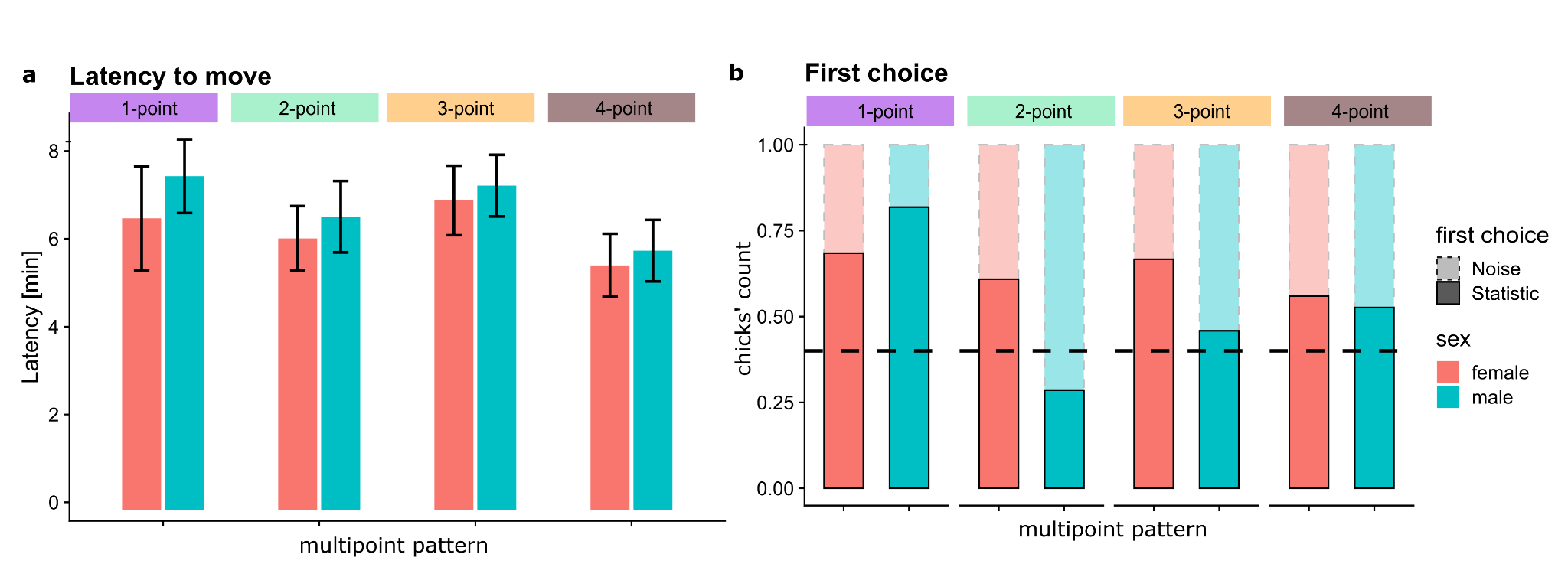

***Supplementary Figure 1.*** Additional data for Experiment 1a. ***a)*** Latency to initiate movement expressed in minutes (mean ± sem). ***b)*** Percentual of chicks’ count showing the first choice for multipoint pattern (shadowed for noise); coloured by sex, dotted horizontal lines indicate the chance level.

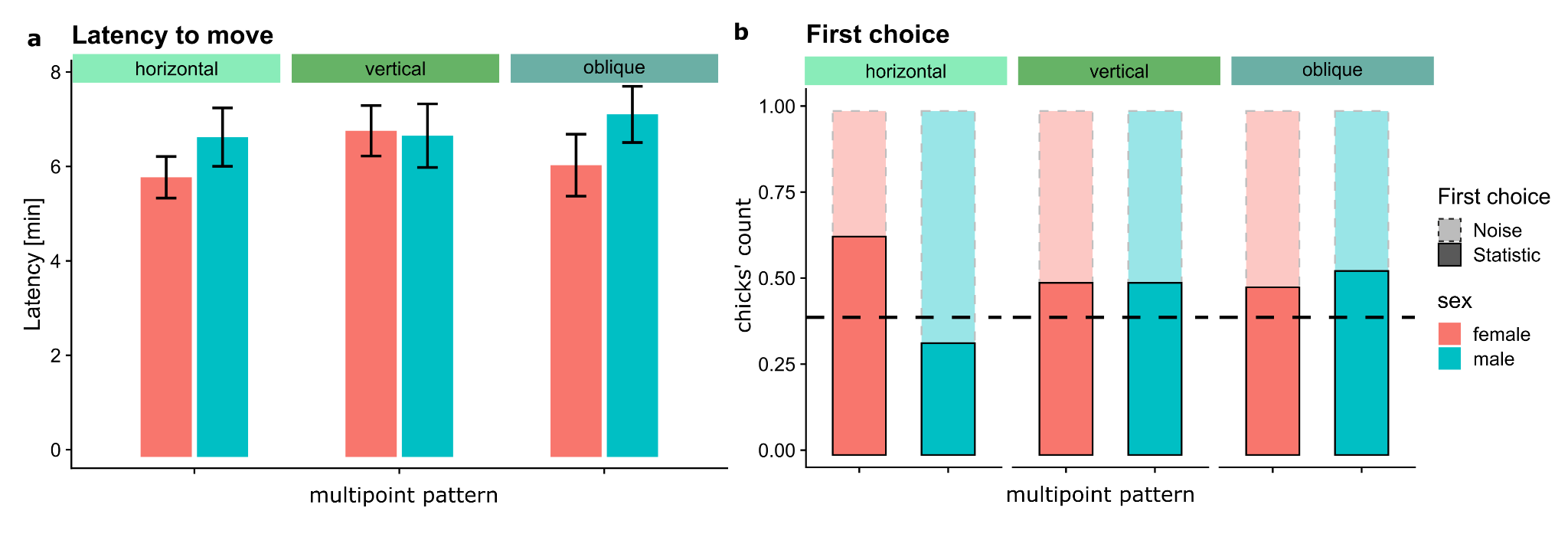

***Supplementary Figure 2.*** Additional data for Experiment 1b. ***a)*** Latency to initiate movement expressed in minutes (mean ± sem). ***b)*** Percentual of chicks’ count showing the first choice for multipoint pattern (shadowed for noise); coloured by sex, dotted horizontal lines indicate the chance level.

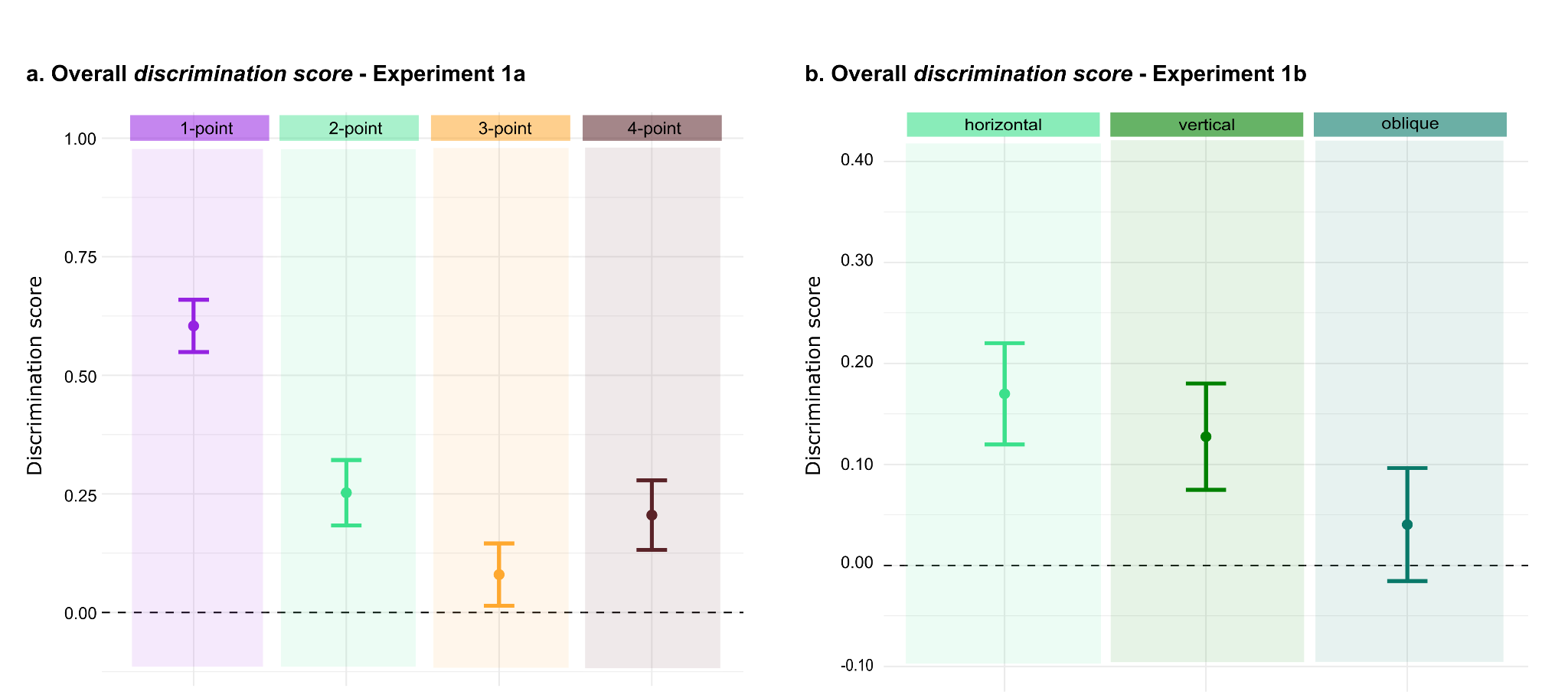

***Supplementary Figure 3.*** Experiment 1, overall discrimination scores averaged across 15 minutes. ***a)*** Overall scores for Experiment 1a. ***b)*** Overall scores for Experiment 1b.

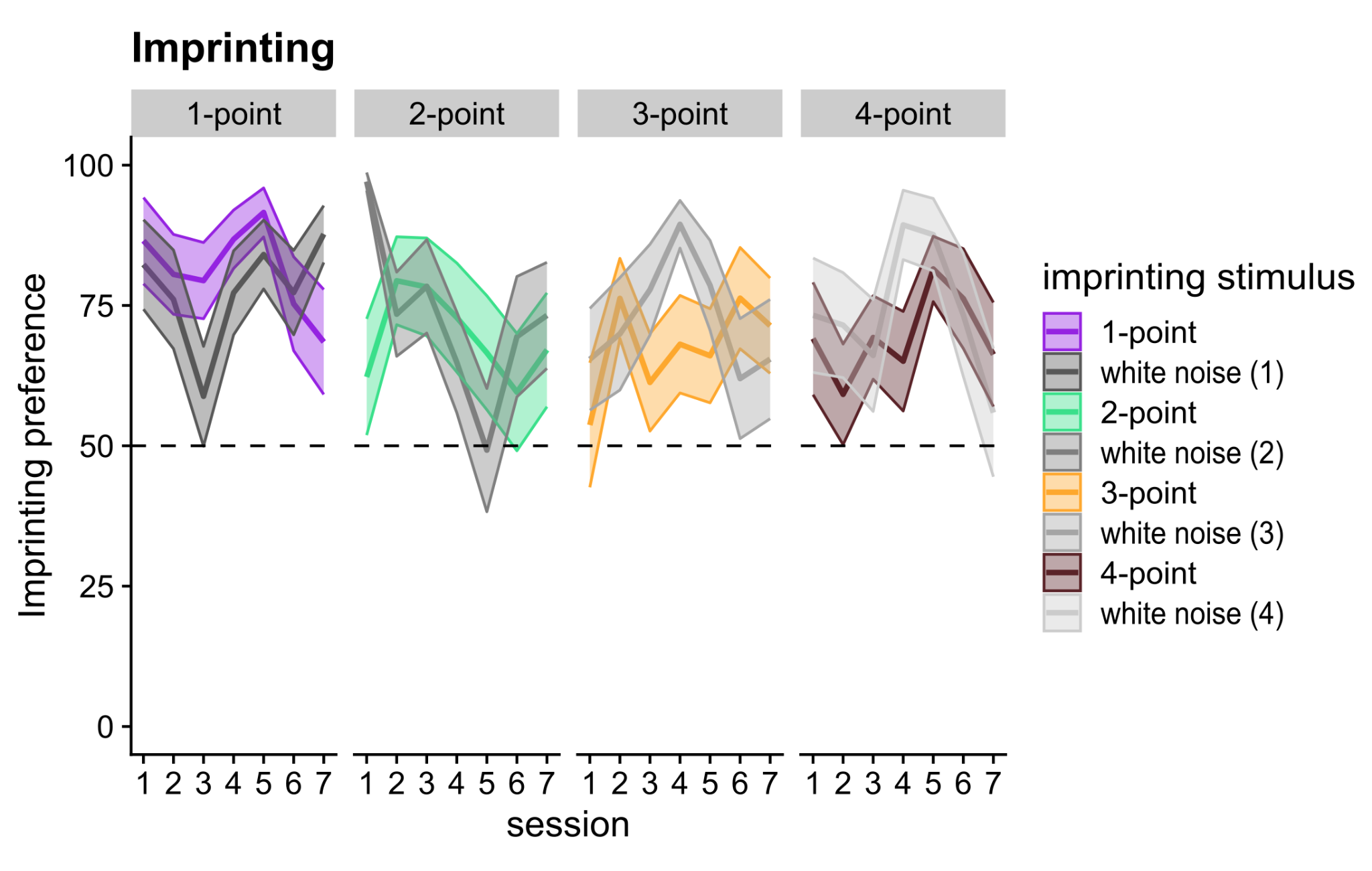

***Supplementary Figure 4.*** Results for Experiment 2, imprinting phase. Average time spent by chicks close to the imprinting stimulus (expressed in percentage, 100% corresponds to 14 hours) during the imprinting phase*.* Curves are coloured by imprinting stimulus (the noise groups are indicated with numbers related to the following test, i.e. (1) if later tested against 1-point etc.); dotted horizontal lines indicate the chance level.

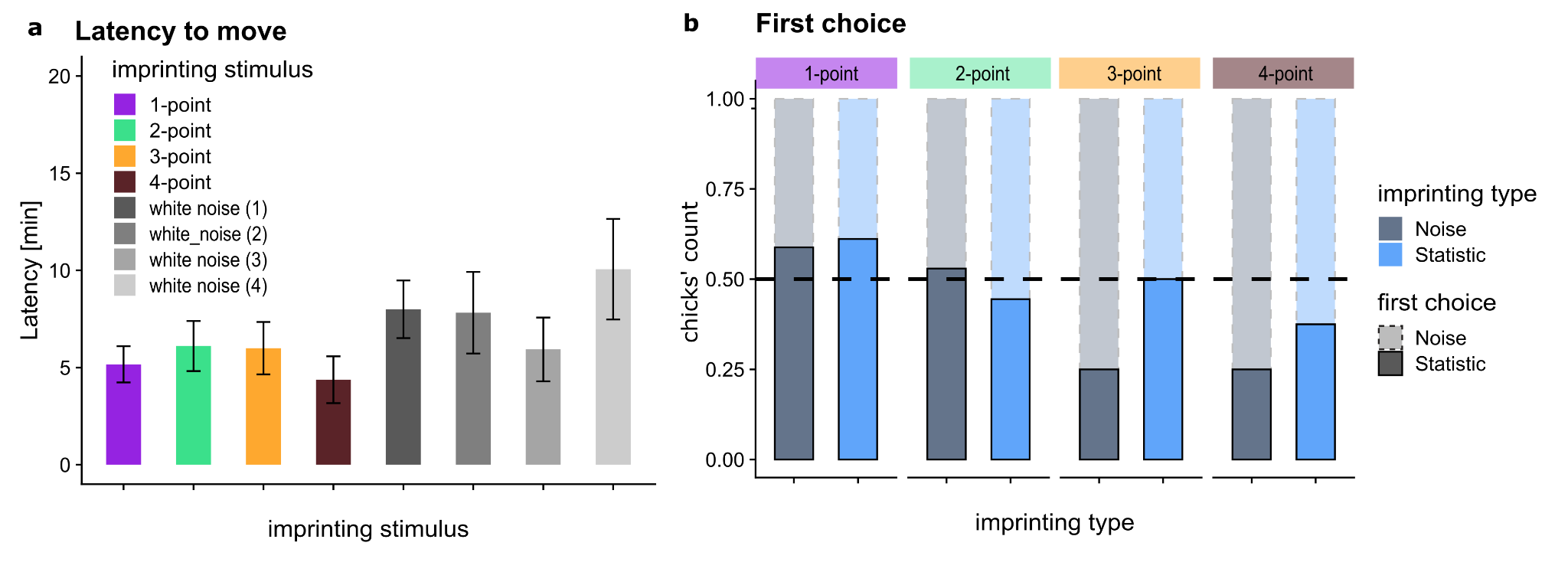

***Supplementary Figure 5.*** Additional data for test phase of Experiment 2. ***a)*** Latency to initiate movement expressed in minutes (mean ± sem). ***b)*** Percentual of chicks’ count showing the first choice for multipoint pattern (shadowed for noise); coloured by type of imprinting (on pattern or on noise), dotted horizontal lines indicate the chance level.

| **1-point** | | | | | | |
| --- | --- | --- | --- | --- | --- | --- |
| **Minute** | **Sex** | **mean PI** | **se PI** | **CI_low** | **CI_high** | **n** |
| 1 | female | 0.31 | 0.21 | -0.13 | 0.76 | 19 |
| 2 | female | 0.25 | 0.20 | -0.18 | 0.68 | 19 |
| 3 | female | 0.35 | 0.19 | -0.04 | 0.75 | 19 |
| 4 | female | 0.35 | 0.19 | -0.04 | 0.75 | 19 |
| 5 | female | 0.56 | 0.17 | 0.20 | 0.93 | 19 |
| 6 | female | 0.51 | 0.19 | 0.11 | 0.91 | 19 |
| 7 | female | 0.51 | 0.19 | 0.11 | 0.91 | 19 |
| 8 | female | 0.72 | 0.15 | 0.41 | 1.03 | 19 |
| 9 | female | 0.71 | 0.15 | 0.40 | 1.02 | 19 |
| 10 | female | 0.65 | 0.15 | 0.32 | 0.97 | 19 |
| 11 | female | 0.68 | 0.15 | 0.37 | 0.99 | 19 |
| 12 | female | 0.75 | 0.14 | 0.45 | 1.05 | 19 |
| 13 | female | 0.75 | 0.14 | 0.45 | 1.05 | 19 |
| 14 | female | 0.50 | 0.18 | 0.12 | 0.88 | 19 |
| 15 | female | 0.39 | 0.20 | -0.04 | 0.82 | 19 |
| 1 | male | 0.61 | 0.17 | 0.26 | 0.95 | 22 |
| 2 | male | 0.69 | 0.13 | 0.41 | 0.96 | 22 |
| 3 | male | 0.53 | 0.15 | 0.22 | 0.84 | 22 |
| 4 | male | 0.66 | 0.14 | 0.38 | 0.95 | 22 |
| 5 | male | 0.66 | 0.14 | 0.38 | 0.94 | 22 |
| 6 | male | 0.69 | 0.11 | 0.46 | 0.93 | 22 |
| 7 | male | 0.63 | 0.14 | 0.34 | 0.91 | 22 |
| 8 | male | 0.71 | 0.13 | 0.43 | 0.98 | 22 |
| 9 | male | 0.72 | 0.11 | 0.48 | 0.95 | 22 |
| 10 | male | 0.71 | 0.11 | 0.47 | 0.94 | 22 |
| 11 | male | 0.52 | 0.17 | 0.16 | 0.87 | 22 |
| 12 | male | 0.68 | 0.12 | 0.43 | 0.93 | 22 |
| 13 | male | 0.74 | 0.11 | 0.51 | 0.98 | 22 |
| 14 | male | 0.76 | 0.13 | 0.49 | 1.02 | 22 |
| 15 | male | 0.68 | 0.15 | 0.37 | 0.98 | 22 |
| **2-point** | | | | | | |
| **Minute** | **Sex** | **mean PI** | **se PI** | **CI_low** | **CI_high** | **n** |
| 1 | female | 0.26 | 0.19 | -0.13 | 0.66 | 23 |
| 2 | female | 0.25 | 0.19 | -0.14 | 0.65 | 23 |
| 3 | female | 0.17 | 0.19 | -0.21 | 0.56 | 23 |
| 4 | female | 0.17 | 0.18 | -0.21 | 0.54 | 23 |
| 5 | female | 0.13 | 0.18 | -0.24 | 0.49 | 23 |
| 6 | female | 0.08 | 0.18 | -0.30 | 0.45 | 23 |
| 7 | female | -0.10 | 0.17 | -0.46 | 0.26 | 23 |
| 8 | female | -0.16 | 0.18 | -0.52 | 0.21 | 23 |
| 9 | female | -0.22 | 0.18 | -0.60 | 0.16 | 23 |
| 10 | female | -0.27 | 0.19 | -0.66 | 0.12 | 23 |
| 11 | female | -0.19 | 0.19 | -0.59 | 0.21 | 23 |
| 12 | female | -0.19 | 0.19 | -0.59 | 0.21 | 23 |
| 13 | female | -0.19 | 0.19 | -0.60 | 0.21 | 23 |
| 14 | female | -0.12 | 0.19 | -0.50 | 0.27 | 23 |
| 15 | female | -0.20 | 0.19 | -0.60 | 0.20 | 23 |
| 1 | male | -0.44 | 0.20 | -0.85 | -0.03 | 21 |
| 2 | male | -0.36 | 0.20 | -0.77 | 0.06 | 21 |
| 3 | male | -0.46 | 0.19 | -0.85 | -0.07 | 21 |
| 4 | male | -0.39 | 0.20 | -0.80 | 0.02 | 21 |
| 5 | male | -0.38 | 0.20 | -0.80 | 0.03 | 21 |
| 6 | male | -0.29 | 0.20 | -0.69 | 0.12 | 21 |
| 7 | male | -0.15 | 0.21 | -0.58 | 0.29 | 21 |
| 8 | male | -0.27 | 0.20 | -0.68 | 0.14 | 21 |
| 9 | male | -0.39 | 0.18 | -0.77 | 0.00 | 21 |
| 10 | male | -0.24 | 0.20 | -0.67 | 0.18 | 21 |
| 11 | male | -0.20 | 0.20 | -0.62 | 0.22 | 21 |
| 12 | male | -0.33 | 0.19 | -0.72 | 0.06 | 21 |
| 13 | male | -0.53 | 0.16 | -0.86 | -0.20 | 21 |
| 14 | male | -0.33 | 0.18 | -0.71 | 0.05 | 21 |
| 15 | male | -0.39 | 0.18 | -0.76 | -0.02 | 21 |
| **3-point** | | | | | | |
| **Minute** | **Sex** | **mean PI** | **se PI** | **CI_low** | **CI_high** | **n** |
| 1 | female | 0.28 | 0.18 | -0.08 | 0.65 | 27 |
| 2 | female | 0.09 | 0.18 | -0.29 | 0.46 | 27 |
| 3 | female | 0.23 | 0.17 | -0.12 | 0.58 | 27 |
| 4 | female | -0.09 | 0.17 | -0.45 | 0.26 | 27 |
| 5 | female | -0.06 | 0.16 | -0.40 | 0.28 | 27 |
| 6 | female | 0.05 | 0.16 | -0.28 | 0.39 | 27 |
| 7 | female | -0.13 | 0.16 | -0.45 | 0.19 | 27 |
| 8 | female | -0.04 | 0.15 | -0.35 | 0.27 | 27 |
| 9 | female | 0.10 | 0.18 | -0.27 | 0.46 | 27 |
| 10 | female | -0.08 | 0.17 | -0.43 | 0.28 | 27 |
| 11 | female | -0.12 | 0.18 | -0.49 | 0.25 | 27 |
| 12 | female | -0.12 | 0.18 | -0.48 | 0.25 | 27 |
| 13 | female | -0.16 | 0.18 | -0.53 | 0.22 | 27 |
| 14 | female | -0.09 | 0.18 | -0.45 | 0.27 | 27 |
| 15 | female | -0.12 | 0.18 | -0.48 | 0.25 | 27 |
| 1 | male | -0.11 | 0.20 | -0.53 | 0.31 | 24 |
| 2 | male | -0.07 | 0.20 | -0.49 | 0.35 | 24 |
| 3 | male | -0.04 | 0.20 | -0.46 | 0.38 | 24 |
| 4 | male | -0.23 | 0.18 | -0.60 | 0.14 | 24 |
| 5 | male | -0.23 | 0.17 | -0.59 | 0.13 | 24 |
| 6 | male | -0.08 | 0.18 | -0.45 | 0.28 | 24 |
| 7 | male | -0.10 | 0.19 | -0.48 | 0.29 | 24 |
| 8 | male | -0.09 | 0.19 | -0.48 | 0.30 | 24 |
| 9 | male | -0.08 | 0.19 | -0.47 | 0.31 | 24 |
| 10 | male | -0.08 | 0.19 | -0.47 | 0.30 | 24 |
| 11 | male | -0.11 | 0.19 | -0.49 | 0.27 | 24 |
| 12 | male | -0.11 | 0.19 | -0.50 | 0.28 | 24 |
| 13 | male | 0.03 | 0.20 | -0.37 | 0.44 | 24 |
| 14 | male | -0.01 | 0.18 | -0.37 | 0.35 | 24 |
| 15 | male | 0.06 | 0.17 | -0.30 | 0.42 | 24 |
| **4-point** | | | | | | |
| **Minute** | **Sex** | **mean PI** | **se PI** | **CI_low** | **CI_high** | **n** |
| 1 | female | 0.07 | 0.19 | -0.33 | 0.47 | 25 |
| 2 | female | 0.11 | 0.19 | -0.28 | 0.50 | 25 |
| 3 | female | 0.04 | 0.18 | -0.34 | 0.42 | 25 |
| 4 | female | 0.11 | 0.19 | -0.28 | 0.50 | 25 |
| 5 | female | 0.21 | 0.19 | -0.17 | 0.60 | 25 |
| 6 | female | 0.26 | 0.17 | -0.10 | 0.62 | 25 |
| 7 | female | 0.30 | 0.18 | -0.08 | 0.68 | 25 |
| 8 | female | 0.30 | 0.17 | -0.06 | 0.66 | 25 |
| 9 | female | 0.35 | 0.18 | -0.02 | 0.72 | 25 |
| 10 | female | 0.38 | 0.18 | 0.01 | 0.75 | 25 |
| 11 | female | 0.31 | 0.18 | -0.06 | 0.67 | 25 |
| 12 | female | 0.39 | 0.18 | 0.02 | 0.76 | 25 |
| 13 | female | 0.35 | 0.18 | -0.02 | 0.72 | 25 |
| 14 | female | 0.35 | 0.18 | -0.02 | 0.72 | 25 |
| 15 | female | 0.39 | 0.18 | 0.02 | 0.75 | 25 |
| 1 | male | 0.00 | 0.23 | -0.47 | 0.48 | 19 |
| 2 | male | -0.01 | 0.23 | -0.49 | 0.46 | 19 |
| 3 | male | -0.07 | 0.23 | -0.55 | 0.42 | 19 |
| 4 | male | -0.01 | 0.22 | -0.49 | 0.46 | 19 |
| 5 | male | 0.09 | 0.21 | -0.36 | 0.53 | 19 |
| 6 | male | 0.04 | 0.22 | -0.42 | 0.51 | 19 |
| 7 | male | 0.32 | 0.20 | -0.10 | 0.74 | 19 |
| 8 | male | 0.32 | 0.20 | -0.10 | 0.74 | 19 |
| 9 | male | 0.23 | 0.21 | -0.21 | 0.67 | 19 |
| 10 | male | 0.03 | 0.22 | -0.43 | 0.49 | 19 |
| 11 | male | 0.11 | 0.20 | -0.31 | 0.53 | 19 |
| 12 | male | 0.25 | 0.21 | -0.19 | 0.68 | 19 |
| 13 | male | 0.26 | 0.20 | -0.15 | 0.68 | 19 |
| 14 | male | 0.29 | 0.20 | -0.13 | 0.71 | 19 |
| 15 | male | 0.13 | 0.21 | -0.31 | 0.57 | 19 |

***Supplementary Table 1.*** Raw reference index in time for Experiment 1a. Average minute by minute of the preference index (PI) for the groups ‘type of multipoint pattern’ and sex. n is the number of chicks per group, se the standard error and CI the 95% confidence interval.

| **horizontal** | | | | | | |
| --- | --- | --- | --- | --- | --- | --- |
| **Minute** | **Sex** | **mean PI** | **se PI** | **CI_low** | **CI_high** | **n** |
| 1 | female | 0.27 | 0.13 | 0.02 | 0.53 | 52 |
| 2 | female | 0.24 | 0.13 | -0.02 | 0.49 | 52 |
| 3 | female | 0.17 | 0.13 | -0.08 | 0.43 | 52 |
| 4 | female | 0.11 | 0.13 | -0.14 | 0.36 | 52 |
| 5 | female | 0.05 | 0.13 | -0.21 | 0.31 | 52 |
| 6 | female | 0.09 | 0.13 | -0.17 | 0.34 | 52 |
| 7 | female | 0.01 | 0.13 | -0.24 | 0.26 | 52 |
| 8 | female | 0.07 | 0.12 | -0.18 | 0.31 | 52 |
| 9 | female | -0.10 | 0.13 | -0.35 | 0.16 | 52 |
| 10 | female | -0.02 | 0.13 | -0.29 | 0.24 | 52 |
| 11 | female | 0.04 | 0.13 | -0.23 | 0.30 | 52 |
| 12 | female | 0.01 | 0.13 | -0.25 | 0.28 | 52 |
| 13 | female | -0.05 | 0.13 | -0.32 | 0.21 | 52 |
| 14 | female | 0.04 | 0.13 | -0.22 | 0.29 | 52 |
| 15 | female | 0.01 | 0.13 | -0.25 | 0.27 | 52 |
| 1 | male | -0.37 | 0.14 | -0.66 | -0.08 | 40 |
| 2 | male | -0.31 | 0.14 | -0.60 | -0.02 | 40 |
| 3 | male | -0.31 | 0.14 | -0.59 | -0.03 | 40 |
| 4 | male | -0.36 | 0.14 | -0.63 | -0.08 | 40 |
| 5 | male | -0.28 | 0.14 | -0.57 | 0.00 | 40 |
| 6 | male | -0.24 | 0.14 | -0.51 | 0.04 | 40 |
| 7 | male | -0.22 | 0.14 | -0.52 | 0.07 | 40 |
| 8 | male | -0.30 | 0.13 | -0.57 | -0.02 | 40 |
| 9 | male | -0.34 | 0.14 | -0.62 | -0.07 | 40 |
| 10 | male | -0.37 | 0.14 | -0.65 | -0.09 | 40 |
| 11 | male | -0.29 | 0.14 | -0.57 | 0.00 | 40 |
| 12 | male | -0.33 | 0.14 | -0.61 | -0.05 | 40 |
| 13 | male | -0.39 | 0.13 | -0.66 | -0.13 | 40 |
| 14 | male | -0.25 | 0.14 | -0.53 | 0.04 | 40 |
| 15 | male | -0.30 | 0.14 | -0.58 | -0.01 | 40 |
| **vertical** | | | | | | |
| **Minute** | **Sex** | **mean PI** | **se PI** | **CI_low** | **CI_high** | **n** |
| 1 | female | 0.02 | 0.15 | -0.27 | 0.31 | 44 |
| 2 | female | 0.00 | 0.14 | -0.28 | 0.28 | 44 |
| 3 | female | 0.06 | 0.13 | -0.21 | 0.33 | 44 |
| 4 | female | 0.03 | 0.14 | -0.25 | 0.32 | 44 |
| 5 | female | 0.07 | 0.14 | -0.21 | 0.35 | 44 |
| 6 | female | 0.08 | 0.15 | -0.22 | 0.38 | 44 |
| 7 | female | 0.05 | 0.14 | -0.24 | 0.33 | 44 |
| 8 | female | -0.01 | 0.14 | -0.29 | 0.27 | 44 |
| 9 | female | 0.13 | 0.14 | -0.15 | 0.40 | 44 |
| 10 | female | 0.17 | 0.13 | -0.10 | 0.44 | 44 |
| 11 | female | 0.16 | 0.14 | -0.12 | 0.43 | 44 |
| 12 | female | 0.04 | 0.14 | -0.25 | 0.32 | 44 |
| 13 | female | 0.08 | 0.14 | -0.19 | 0.36 | 44 |
| 14 | female | 0.03 | 0.14 | -0.25 | 0.31 | 44 |
| 15 | female | -0.05 | 0.14 | -0.34 | 0.23 | 44 |
| 1 | male | -0.01 | 0.16 | -0.32 | 0.31 | 36 |
| 2 | male | 0.11 | 0.15 | -0.20 | 0.41 | 36 |
| 3 | male | 0.14 | 0.15 | -0.16 | 0.43 | 36 |
| 4 | male | 0.17 | 0.15 | -0.14 | 0.47 | 36 |
| 5 | male | 0.15 | 0.15 | -0.16 | 0.45 | 36 |
| 6 | male | 0.27 | 0.15 | -0.03 | 0.58 | 36 |
| 7 | male | 0.38 | 0.14 | 0.10 | 0.67 | 36 |
| 8 | male | 0.37 | 0.14 | 0.09 | 0.66 | 36 |
| 9 | male | 0.34 | 0.14 | 0.05 | 0.64 | 36 |
| 10 | male | 0.20 | 0.15 | -0.11 | 0.50 | 36 |
| 11 | male | 0.21 | 0.15 | -0.09 | 0.50 | 36 |
| 12 | male | 0.20 | 0.15 | -0.11 | 0.50 | 36 |
| 13 | male | 0.25 | 0.15 | -0.05 | 0.55 | 36 |
| 14 | male | 0.28 | 0.15 | -0.02 | 0.58 | 36 |
| 15 | male | 0.18 | 0.15 | -0.12 | 0.48 | 36 |
| **oblique** | | | | | | |
| **Minute** | **Sex** | **mean PI** | **se PI** | **CI_low** | **CI_high** | **n** |
| 1 | female | -0.04 | 0.16 | -0.35 | 0.28 | 39 |
| 2 | female | -0.06 | 0.15 | -0.37 | 0.25 | 39 |
| 3 | female | 0.02 | 0.15 | -0.28 | 0.31 | 39 |
| 4 | female | -0.01 | 0.15 | -0.32 | 0.29 | 39 |
| 5 | female | 0.11 | 0.15 | -0.19 | 0.41 | 39 |
| 6 | female | 0.07 | 0.15 | -0.23 | 0.37 | 39 |
| 7 | female | -0.05 | 0.15 | -0.35 | 0.25 | 39 |
| 8 | female | -0.07 | 0.16 | -0.39 | 0.24 | 39 |
| 9 | female | -0.08 | 0.15 | -0.39 | 0.23 | 39 |
| 10 | female | -0.03 | 0.16 | -0.35 | 0.28 | 39 |
| 11 | female | 0.00 | 0.15 | -0.31 | 0.31 | 39 |
| 12 | female | 0.01 | 0.15 | -0.29 | 0.31 | 39 |
| 13 | female | 0.04 | 0.15 | -0.27 | 0.36 | 39 |
| 14 | female | -0.01 | 0.16 | -0.33 | 0.31 | 39 |
| 15 | female | -0.01 | 0.16 | -0.33 | 0.31 | 39 |
| 1 | male | 0.05 | 0.15 | -0.26 | 0.35 | 43 |
| 2 | male | 0.09 | 0.14 | -0.19 | 0.38 | 43 |
| 3 | male | 0.07 | 0.15 | -0.23 | 0.37 | 43 |
| 4 | male | 0.10 | 0.15 | -0.20 | 0.40 | 43 |
| 5 | male | 0.06 | 0.15 | -0.24 | 0.36 | 43 |
| 6 | male | -0.01 | 0.14 | -0.30 | 0.28 | 43 |
| 7 | male | 0.12 | 0.14 | -0.16 | 0.40 | 43 |
| 8 | male | 0.11 | 0.14 | -0.17 | 0.39 | 43 |
| 9 | male | 0.17 | 0.14 | -0.12 | 0.46 | 43 |
| 10 | male | 0.06 | 0.14 | -0.22 | 0.35 | 43 |
| 11 | male | 0.05 | 0.15 | -0.25 | 0.35 | 43 |
| 12 | male | 0.06 | 0.15 | -0.23 | 0.35 | 43 |
| 13 | male | -0.07 | 0.15 | -0.36 | 0.23 | 43 |
| 14 | male | 0.01 | 0.14 | -0.28 | 0.30 | 43 |
| 15 | male | 0.09 | 0.14 | -0.20 | 0.37 | 43 |

***Supplementary Table 2.*** Raw reference index in time for Experiment 1b. Average minute by minute of the preference index (PI) for the groups ‘type of multipoint pattern’ and sex. n is the number of chicks per group, se the standard error and CI the 95% confidence interval.

| **1-point** | | | | | | |
| --- | --- | --- | --- | --- | --- | --- |
| **Minute** | **Imprinted on** | **mean PI** | **se PI** | **CI_low** | **CI_high** | **n** |
| 1 | Noise | 0.20 | 0.24 | -0.31 | 0.71 | 17 |
| 2 | Noise | 0.12 | 0.24 | -0.38 | 0.63 | 17 |
| 3 | Noise | 0.12 | 0.24 | -0.38 | 0.63 | 17 |
| 4 | Noise | 0.15 | 0.23 | -0.34 | 0.64 | 17 |
| 5 | Noise | 0.18 | 0.24 | -0.34 | 0.70 | 17 |
| 6 | Noise | 0.15 | 0.24 | -0.35 | 0.66 | 17 |
| 7 | Noise | -0.11 | 0.24 | -0.61 | 0.40 | 17 |
| 8 | Noise | 0.07 | 0.25 | -0.46 | 0.59 | 17 |
| 9 | Noise | 0.07 | 0.23 | -0.42 | 0.56 | 17 |
| 10 | Noise | 0.01 | 0.24 | -0.50 | 0.51 | 17 |
| 11 | Noise | 0.01 | 0.24 | -0.50 | 0.52 | 17 |
| 12 | Noise | 0.11 | 0.24 | -0.39 | 0.61 | 17 |
| 13 | Noise | -0.05 | 0.23 | -0.54 | 0.44 | 17 |
| 14 | Noise | -0.05 | 0.23 | -0.54 | 0.44 | 17 |
| 15 | Noise | -0.15 | 0.21 | -0.60 | 0.30 | 17 |
| 1 | Statistic | 0.17 | 0.23 | -0.31 | 0.65 | 18 |
| 2 | Statistic | 0.26 | 0.22 | -0.21 | 0.74 | 18 |
| 3 | Statistic | 0.15 | 0.21 | -0.30 | 0.61 | 18 |
| 4 | Statistic | 0.08 | 0.21 | -0.36 | 0.52 | 18 |
| 5 | Statistic | 0.05 | 0.22 | -0.41 | 0.51 | 18 |
| 6 | Statistic | 0.05 | 0.22 | -0.41 | 0.51 | 18 |
| 7 | Statistic | 0.10 | 0.22 | -0.37 | 0.58 | 18 |
| 8 | Statistic | 0.05 | 0.22 | -0.41 | 0.51 | 18 |
| 9 | Statistic | 0.03 | 0.23 | -0.45 | 0.52 | 18 |
| 10 | Statistic | 0.26 | 0.19 | -0.14 | 0.66 | 18 |
| 11 | Statistic | 0.35 | 0.21 | -0.09 | 0.79 | 18 |
| 12 | Statistic | 0.35 | 0.21 | -0.09 | 0.79 | 18 |
| 13 | Statistic | 0.27 | 0.21 | -0.17 | 0.70 | 18 |
| 14 | Statistic | 0.26 | 0.21 | -0.18 | 0.70 | 18 |
| 15 | Statistic | 0.32 | 0.21 | -0.12 | 0.77 | 18 |
| **2-point** | | | | | | |
| **Minute** | **Imprinted on** | **mean PI** | **se PI** | **CI_low** | **CI_high** | **n** |
| 1 | Noise | 0.10 | 0.24 | -0.41 | 0.61 | 17 |
| 2 | Noise | 0.07 | 0.23 | -0.43 | 0.56 | 17 |
| 3 | Noise | 0.01 | 0.22 | -0.47 | 0.48 | 17 |
| 4 | Noise | 0.01 | 0.22 | -0.47 | 0.48 | 17 |
| 5 | Noise | 0.01 | 0.22 | -0.47 | 0.48 | 17 |
| 6 | Noise | -0.11 | 0.21 | -0.55 | 0.33 | 17 |
| 7 | Noise | -0.17 | 0.21 | -0.62 | 0.28 | 17 |
| 8 | Noise | -0.11 | 0.22 | -0.58 | 0.36 | 17 |
| 9 | Noise | 0.01 | 0.22 | -0.46 | 0.49 | 17 |
| 10 | Noise | -0.11 | 0.22 | -0.59 | 0.36 | 17 |
| 11 | Noise | -0.11 | 0.22 | -0.58 | 0.37 | 17 |
| 12 | Noise | 0.00 | 0.23 | -0.47 | 0.48 | 17 |
| 13 | Noise | 0.01 | 0.22 | -0.47 | 0.48 | 17 |
| 14 | Noise | -0.04 | 0.23 | -0.53 | 0.44 | 17 |
| 15 | Noise | 0.01 | 0.23 | -0.47 | 0.48 | 17 |
| 1 | Statistic | -0.17 | 0.23 | -0.65 | 0.31 | 18 |
| 2 | Statistic | -0.12 | 0.24 | -0.62 | 0.38 | 18 |
| 3 | Statistic | 0.00 | 0.24 | -0.49 | 0.50 | 18 |
| 4 | Statistic | 0.04 | 0.23 | -0.45 | 0.53 | 18 |
| 5 | Statistic | 0.05 | 0.23 | -0.44 | 0.54 | 18 |
| 6 | Statistic | 0.05 | 0.23 | -0.44 | 0.54 | 18 |
| 7 | Statistic | 0.05 | 0.23 | -0.45 | 0.54 | 18 |
| 8 | Statistic | 0.09 | 0.23 | -0.39 | 0.57 | 18 |
| 9 | Statistic | 0.10 | 0.22 | -0.37 | 0.58 | 18 |
| 10 | Statistic | 0.06 | 0.23 | -0.43 | 0.55 | 18 |
| 11 | Statistic | 0.16 | 0.23 | -0.33 | 0.64 | 18 |
| 12 | Statistic | 0.05 | 0.23 | -0.44 | 0.54 | 18 |
| 13 | Statistic | 0.17 | 0.23 | -0.31 | 0.65 | 18 |
| 14 | Statistic | 0.17 | 0.22 | -0.30 | 0.64 | 18 |
| 15 | Statistic | 0.27 | 0.22 | -0.20 | 0.74 | 18 |
| **3-point** | | | | | | |
| **Minute** | **Imprinted on** | **mean PI** | **se PI** | **CI_low** | **CI_high** | **n** |
| 1 | Noise | -0.44 | 0.22 | -0.91 | 0.03 | 16 |
| 2 | Noise | -0.51 | 0.20 | -0.93 | -0.08 | 16 |
| 3 | Noise | -0.52 | 0.20 | -0.95 | -0.10 | 16 |
| 4 | Noise | -0.50 | 0.20 | -0.92 | -0.07 | 16 |
| 5 | Noise | -0.42 | 0.22 | -0.89 | 0.05 | 16 |
| 6 | Noise | -0.30 | 0.22 | -0.76 | 0.16 | 16 |
| 7 | Noise | -0.30 | 0.22 | -0.76 | 0.16 | 16 |
| 8 | Noise | -0.36 | 0.22 | -0.82 | 0.11 | 16 |
| 9 | Noise | -0.36 | 0.22 | -0.83 | 0.11 | 16 |
| 10 | Noise | -0.37 | 0.22 | -0.83 | 0.10 | 16 |
| 11 | Noise | -0.36 | 0.22 | -0.83 | 0.10 | 16 |
| 12 | Noise | -0.36 | 0.22 | -0.83 | 0.10 | 16 |
| 13 | Noise | -0.36 | 0.22 | -0.83 | 0.11 | 16 |
| 14 | Noise | -0.30 | 0.22 | -0.76 | 0.16 | 16 |
| 15 | Noise | -0.26 | 0.21 | -0.72 | 0.19 | 16 |
| 1 | Statistic | -0.02 | 0.24 | -0.52 | 0.48 | 18 |
| 2 | Statistic | -0.19 | 0.23 | -0.67 | 0.28 | 18 |
| 3 | Statistic | -0.28 | 0.21 | -0.72 | 0.16 | 18 |
| 4 | Statistic | -0.19 | 0.21 | -0.63 | 0.25 | 18 |
| 5 | Statistic | -0.19 | 0.21 | -0.64 | 0.25 | 18 |
| 6 | Statistic | -0.22 | 0.21 | -0.66 | 0.21 | 18 |
| 7 | Statistic | -0.19 | 0.21 | -0.64 | 0.26 | 18 |
| 8 | Statistic | -0.29 | 0.22 | -0.76 | 0.19 | 18 |
| 9 | Statistic | -0.29 | 0.22 | -0.76 | 0.18 | 18 |
| 10 | Statistic | -0.17 | 0.23 | -0.66 | 0.32 | 18 |
| 11 | Statistic | -0.27 | 0.22 | -0.74 | 0.19 | 18 |
| 12 | Statistic | -0.28 | 0.22 | -0.76 | 0.19 | 18 |
| 13 | Statistic | -0.29 | 0.22 | -0.76 | 0.18 | 18 |
| 14 | Statistic | -0.27 | 0.22 | -0.74 | 0.19 | 18 |
| 15 | Statistic | -0.28 | 0.22 | -0.76 | 0.19 | 18 |
| **4-point** | | | | | | |
| **Minute** | **Imprinted on** | **mean PI** | **se PI** | **CI_low** | **CI_high** | **n** |
| 1 | Noise | -0.46 | 0.22 | -0.93 | 0.00 | 16 |
| 2 | Noise | -0.44 | 0.20 | -0.86 | -0.01 | 16 |
| 3 | Noise | -0.36 | 0.22 | -0.83 | 0.10 | 16 |
| 4 | Noise | -0.48 | 0.20 | -0.91 | -0.05 | 16 |
| 5 | Noise | -0.49 | 0.20 | -0.92 | -0.06 | 16 |
| 6 | Noise | -0.48 | 0.20 | -0.91 | -0.05 | 16 |
| 7 | Noise | -0.49 | 0.20 | -0.91 | -0.06 | 16 |
| 8 | Noise | -0.42 | 0.22 | -0.90 | 0.05 | 16 |
| 9 | Noise | -0.55 | 0.20 | -0.98 | -0.12 | 16 |
| 10 | Noise | -0.54 | 0.20 | -0.97 | -0.11 | 16 |
| 11 | Noise | -0.49 | 0.20 | -0.92 | -0.06 | 16 |
| 12 | Noise | -0.49 | 0.20 | -0.92 | -0.06 | 16 |
| 13 | Noise | -0.49 | 0.20 | -0.92 | -0.06 | 16 |
| 14 | Noise | -0.54 | 0.20 | -0.97 | -0.12 | 16 |
| 15 | Noise | -0.55 | 0.20 | -0.98 | -0.12 | 16 |
| 1 | Statistic | -0.27 | 0.24 | -0.79 | 0.24 | 16 |
| 2 | Statistic | -0.33 | 0.23 | -0.82 | 0.17 | 16 |
| 3 | Statistic | -0.38 | 0.24 | -0.89 | 0.12 | 16 |
| 4 | Statistic | -0.38 | 0.24 | -0.89 | 0.12 | 16 |
| 5 | Statistic | -0.38 | 0.24 | -0.89 | 0.12 | 16 |
| 6 | Statistic | -0.38 | 0.24 | -0.89 | 0.12 | 16 |
| 7 | Statistic | -0.38 | 0.24 | -0.89 | 0.12 | 16 |
| 8 | Statistic | -0.38 | 0.24 | -0.89 | 0.12 | 16 |
| 9 | Statistic | -0.38 | 0.24 | -0.89 | 0.12 | 16 |
| 10 | Statistic | -0.38 | 0.24 | -0.89 | 0.12 | 16 |
| 11 | Statistic | -0.38 | 0.24 | -0.89 | 0.12 | 16 |
| 12 | Statistic | -0.38 | 0.24 | -0.89 | 0.12 | 16 |
| 13 | Statistic | -0.38 | 0.24 | -0.89 | 0.12 | 16 |
| 14 | Statistic | -0.38 | 0.24 | -0.89 | 0.12 | 16 |
| 15 | Statistic | -0.38 | 0.24 | -0.89 | 0.12 | 16 |

***Supplementary Table 3.*** Raw reference index in time for Experiment 2 - test phase. Average minute by minute of the preference index (PI) for the groups ‘type of multipoint pattern’ and ‘type of imprinting’. n is the number of chicks per group, se the standard error and CI the 95% confidence interval.
